# Structural and functional basis of VLDLR receptor usage by Eastern equine encephalitis virus

**DOI:** 10.1101/2023.11.15.567188

**Authors:** Lucas J. Adams, Saravanan Raju, Hongming Ma, Theron Gilliland, Douglas S. Reed, William B. Klimstra, Daved H. Fremont, Michael S. Diamond

## Abstract

The very low-density lipoprotein receptor (VLDLR) is comprised of eight LDLR type A (LA) domains and supports entry of distantly related Eastern equine encephalitis (EEEV) and Semliki Forest (SFV) alphaviruses. Here, by resolving multiple cryo-electron microscopy structures of EEEV-VLDLR complexes and performing mutagenesis and functional studies, we show that EEEV uses multiple sites (E1/E2 cleft and E2 A domain) to engage different LA domains simultaneously. However, no single LA domain is necessary or sufficient to support efficient EEEV infection, highlighting complexity in domain usage. Whereas all EEEV strains show conservation of two VLDLR binding sites, the EEEV PE–6 strain and other EEE complex members feature a single amino acid substitution that mediates binding of LA domains to an additional site on the E2 B domain. These structural and functional analyses informed the design of a minimal VLDLR decoy receptor that neutralizes EEEV infection and protects mice from lethal challenge.

## INTRODUCTION

Alphaviruses are arthropod-transmitted, single-stranded positive-sense RNA viruses of the *Togaviridae* family. These enveloped viruses infect a range of vertebrate hosts, including humans, non-human primates, horses, rodents, and birds, and have been categorized as “Old World” or “New World” based on geographic origins.^1,2^ Old World alphaviruses include Chikungunya (CHIKV), Ross River (RRV), Mayaro (MAYV), O’nyong-nyong (ONNV), and Semliki Forest (SFV) viruses, many of which generally cause acute debilitating arthritis and chronic arthralgia in some patients.^3^ New World alphaviruses include Eastern (EEEV), Venezuelan (VEEV), and Western (WEEV) equine encephalitis viruses, which cause neurological disease with high rates of morbidity, mortality, and neurological sequelae.^4^ EEEV, the most pathogenic of the encephalitic alphaviruses, causes sporadic outbreaks across North America, with case fatality rates exceeding 30% in hospitalized patients.^5,6^ Though naturally transmitted by mosquitoes, encephalitic alphaviruses also can be spread via aerosolization and have been weaponized in the past.^7–11^ At present, there are no approved countermeasures for any alphavirus infection.^12^

The mature alphavirus virion is ∼70 nm in diameter and composed of glycoproteins E1 and E2, which dimerize, and an internal capsid protein that packages the RNA genome. The virion exhibits *T* = 4 icosahedral symmetry with 240 E1/E2 heterodimers arranged as 80 trimeric spikes on icosahedral three-fold (*i3*, n = 20) and quasi-threefold (*q3*, n = 60) axes of symmetry. Each asymmetric unit (n = 60) consists of a complete *q3* trimer and a single *i3* E1/E2 heterodimer.^13,14^ Within each heterodimer, the E2 glycoprotein is preferentially exposed and shields the majority of the E1 glycoprotein from solvent, including the conserved fusion loop required for escape from the endosome and nucleocapsid penetration into the cytoplasm.^15–17^ The B domain of E2 is most distal from the viral membrane, forming the three vertices of the trimeric spike, and the E2 A domain is positioned at the axial interface of the trimer. The E2 A and B domains are the principal targets of neutralizing antibodies.^18–26^

A key initial step in the alphavirus infection cycle is engagement with host receptors that promote attachment and entry into cells.^27^ While virus binding to cells is assisted by interactions with attachment factors such as heparan sulfate proteoglycans,^28–36^ specific, proteinaceous receptors enable efficient cell entry. For example, the immunoglobulin (Ig)-like domain containing molecule MXRA8 is a receptor for several members of the Semliki forest virus (SFV) complex (*e.g*., CHIKV, RRV, MAYV, and ONNV),^37–39^ and the low-density lipoprotein receptor (LDLR)-related family member LDLRAD3 is a receptor specifically for VEEV.^40^ Cryo-electron microscopy (cryo-EM) reconstructions of MXRA8 bound to CHIKV and LDLRAD3 bound to VEEV revealed that these unrelated receptors bind their respective viruses within an analogous “cleft” formed by neighboring E1/E2 heterodimers within the trimeric spike.^41–44^ As MXRA8 and LDLRAD3 are not used by all alphaviruses, the existence of additional receptors for other family members was postulated. Indeed, a recent study identified the related proteins VLDLR and ApoER2 (encoded by *LRP8)* as receptors for EEEV, SFV, and to a lesser extent for Sindbis virus (SINV).^45^ For VLDLR, the ligand binding domain (LBD) comprised of eight LDLR type A (LA) repeats was necessary and sufficient to mediate infection, and the interaction of VLDLR LA domains with a single site on SFV E1 domain III was recently reported.^46^ However, the mechanism by which EEEV engages VLDLR has not been determined.

Here, we characterize multiple cryo-EM structures of full-length VLDLR and VLDLR fragments in complex with EEEV virus-like particles (VLPs). This analysis reveals a distinct mode of interaction compared to SFV. For EEEV strain PE–6, multiple different VLDLR LA domains can bind to three distinct sites on the virus: a cleft formed by E1/E2 heterodimers, a shelf on the E2 A domain, and a separate site on the lateral surface of the E2 B domain. Any one of these three LA domain-binding sites is dispensable for efficient, VLDLR-dependent infection by EEEV PE–6. Although critical basic residues mediating interaction with VLDLR in the E1/E2 cleft and the E2 A domain site are conserved across all analyzed EEEV strains, the B domain site, which uses residue K206, is uniquely present in EEEV PE–6 and a few other EEE complex strains, enabling enhanced affinity and versatility of VLDLR domain usage. Together, our data establish the principal modes of VLDLR LA domain engagement with different binding sites on EEEV. Using this knowledge, we developed a VLDLR LA(1–2)-Fc decoy receptor that neutralizes EEEV infection and protects mice against lethal EEEV challenge by an aerosol route.

## RESULTS

### Structure of EEEV in complex with full-length VLDLR

The ligand binding ectodomain (LBD) of VLDLR features eight cysteine-rich LDLR class A (LA) repeats and is necessary and sufficient to mediate infection by EEEV.^45^ To elucidate the structural basis for recognition of VLDLR by EEEV, we reconstructed EEEV VLPs (strain PE–6) alone or in complex with full-length VLDLR via cryo-EM (**Fig S1, S2A-B, and Table S1**), achieving respective resolutions of 3.78 Å and 4.75 Å with icosahedral symmetry imposed. In agreement with previous EEEV VLP or virus structures,^47,48^ the EEEV VLP exhibited *T* = 4 icosahedral symmetry with 80 trimeric spikes composed of E1/E2 heterodimers (**Fig 1A**, *top panel*). Additional densities consistent in size with LA repeats were evident in the VLDLR-bound structure (**Fig 1A**, *bottom panel*). To orient the VLDLR LA domains within these densities, we performed focused refinement of the asymmetric unit (ASU), generating apo and bound structures at 2.86- and 3.89-Å resolution, respectively (**Fig 1B**).

**Figure 1.**
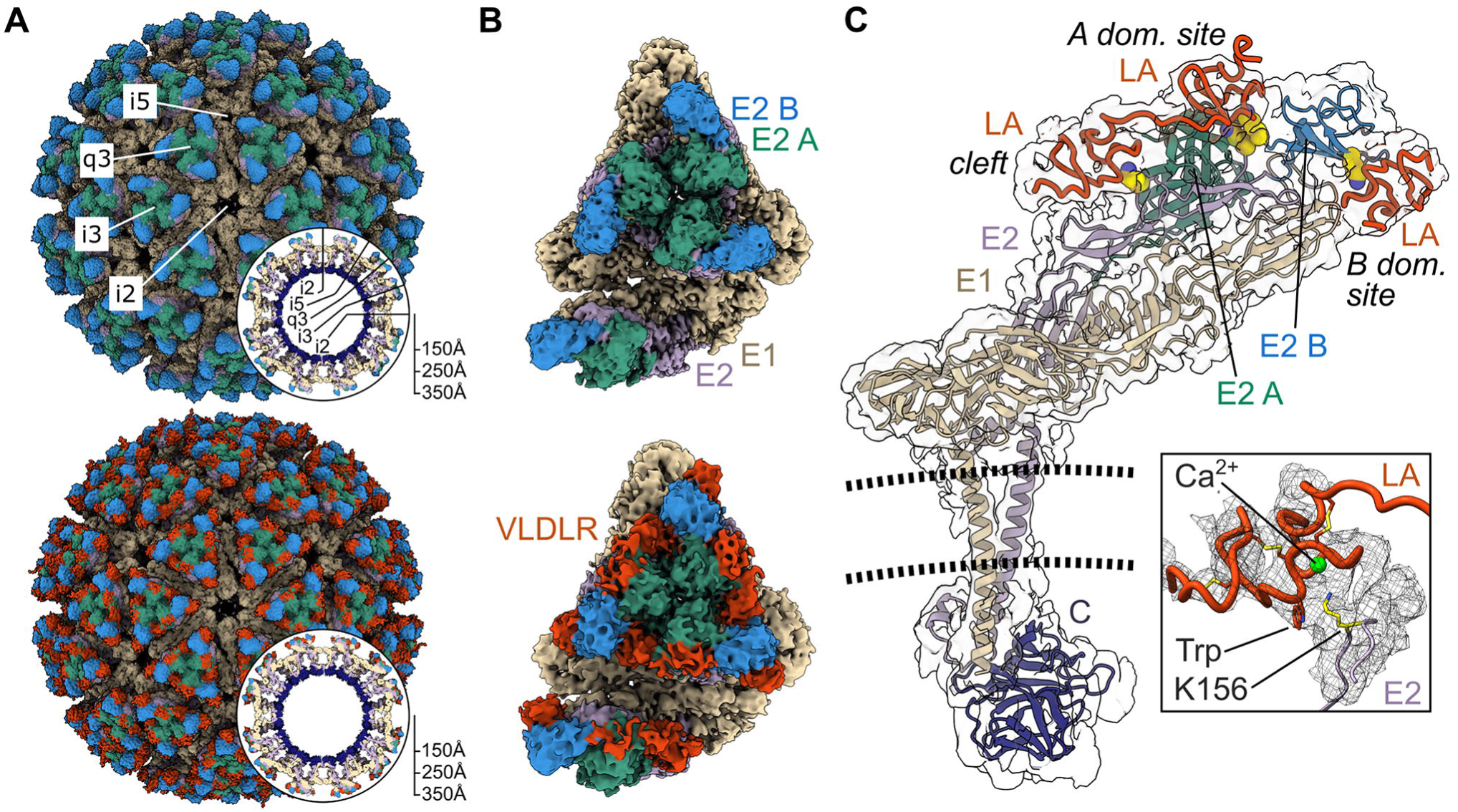
Cryo-EM structure of EEEV PE–6 in complex with VLDLR. (**A**) Icosahedral reconstructions of EEEV PE–6 VLP alone (*upper panel*) or in complex with full-length VLDLR (*lower panel*) with 2-fold (i2), 5-fold (i5), 3-fold (i3), and quasi-3-fold (q3) axes designated. Central sections are shown in round insets. Proteins are differentially colored with E1 in tan, E2 A domain in sea green, E2 B domain in blue, the remainder of E2 colored purple, and VLDLR shown in orange. (**B**) Focused reconstructions of the EEEV asymmetric unit alone (*upper panel*) or in complex with full-length VLDLR (*lower panel*), colored as in (**A**). (**C**) An atomic model of a single E1/E2 heterodimer with non-descript LA domains docked and morphed into the experimental electron density map, colored as in (**A**) with capsid shown in navy and the lipid bilayer depicted as dashed lines. Interfacial lysines are highlighted in yellow, with the region near K156 magnified in the inset.

The ASU of the EEEV-VLDLR complex featured connected densities in the cleft formed between E1/E2 heterodimers and on the slanted shelf of the E2 A domain, as well as a third, lower-resolution density on the outward-facing side of the E2 B domain and E1 domain II. Although this structure did not allow unambiguous assignment of specific LA domains, each LA domain was oriented clearly with conserved calcium-coordination sites proximal to residues K156 (cleft), K231/K232 (A domain), and K206 (B domain) of EEEV E2 (**Fig 1C**). This reconstruction suggests a binding strategy analogous to that observed for SFV and VLDLR, in that a basic residue (K345 of SFV E1) engages a conserved aromatic residue (*e.g*., tryptophan [Trp]) and negatively-charged, calcium-coordinating residues of the LA domain.^46^ However, in contrast to SFV, which engages VLDLR via a single site on E1 domain III, our structure suggests that EEEV uses three distinct sites on the viral glycoproteins for concomitant and/or redundant use of VLDLR LA domains to mediate binding. LA domain densities were not seen near domain III of EEEV E1.

### Mapping of LA domains recognized by EEEV

To assess the role of each LA domain in mediating EEEV infection, we generated domain-swapped variants of VLDLR with the LA1 domain of the related molecule, LDLRAD3, a receptor for VEEV (**Fig 2A and S3A**) based on an approach used to map receptor domain usage of adeno-associated virus.^49^ Importantly, the LA1 domain of LDLRAD3 does not support infection by EEEV.^40,45^ We transduced K562 erythroleukemic cells, which lack endogenous VLDLR expression and are non-permissive to EEEV (**Fig S3B**), with N-terminally Flag-tagged constructs in which a single LA repeat of VLDLR was replaced with LDLRAD3-LA1 and then confirmed cell surface expression by flow cytometry (**Fig S3C**). To assess infection at a lower biosafety containment level, we used an established Sindbis virus (SINV) chimeric GFP reporter virus, in which the structural genes of SINV are replaced with those of EEEV (SINV-EEEV; PE–6 strain) (**Fig S3D**). Unexpectedly, infection was supported by all chimeric constructs, suggesting functional redundancy of at least some of the VLDLR LA domains (**Fig 2B**). As an additional control, we replaced the LBD of VLDLR with LDLRAD3 (LA1–LA3); this chimera did not support infection of K562 cells by EEEV (**Fig 2B**) but did promote VEEV infection (**Fig S3E**), which indicates that LDLRAD3 LA domains are unable to functionally “stand-in” for VLDLR LA domains. These results suggest multiple LA domains can bind to EEEV in a redundant fashion and/or no single site is necessary to support infection.

**Figure 2.**
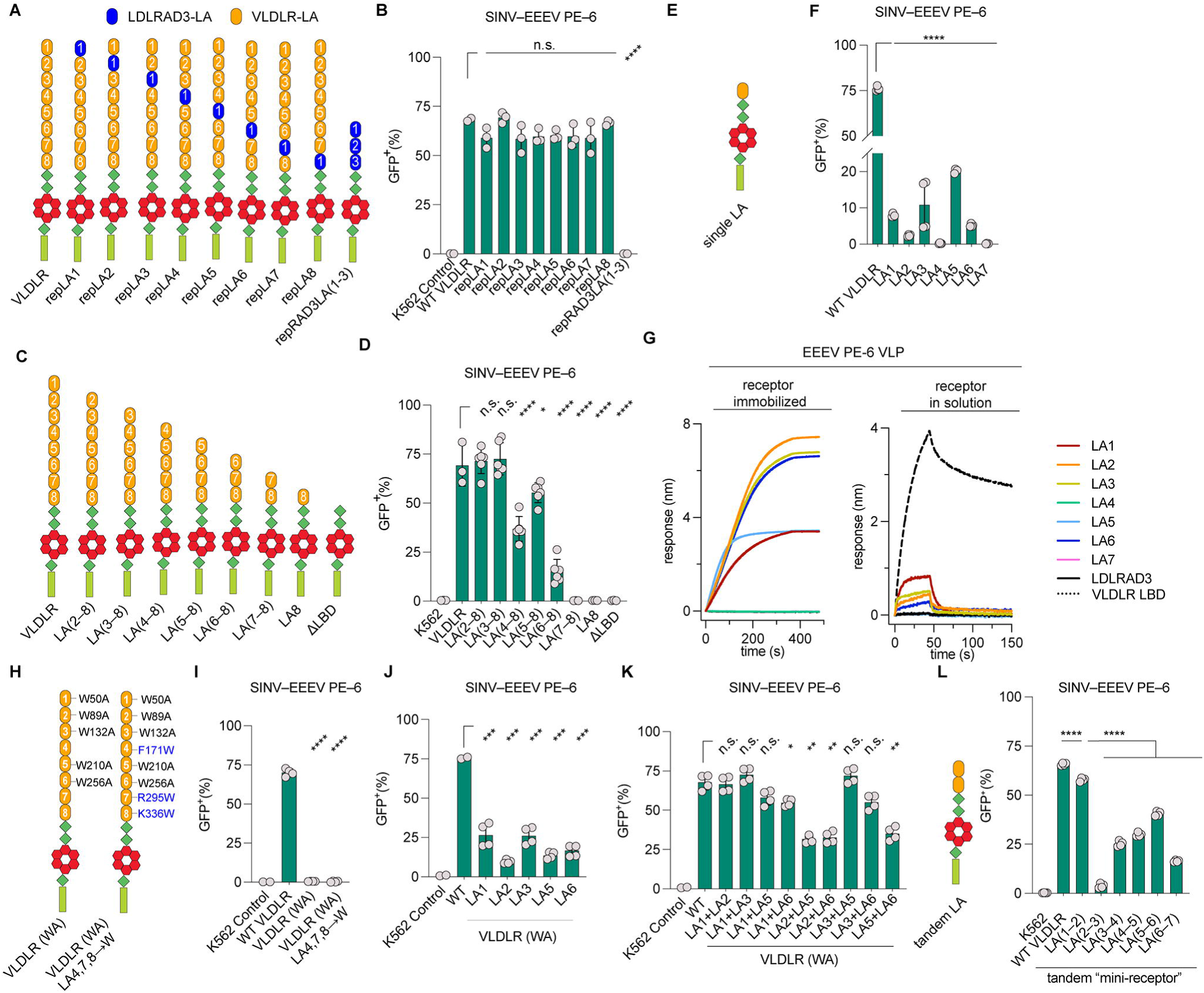
No single LA domain of the VLDLR LBD is required to support EEEV. (**A**) Scheme of LDLRAD3-LA1 domain replacement of VLDLR LA domains. (**B, D, F, H**) SINV-EEEV-GFP PE–6 infection of K562 cells transduced with indicated N-terminal Flag-tagged constructs as quantified by GFP expression using flow cytometry. (**C**) Scheme of N-terminal VLDLR LA domain truncation constructs. (**E**) Scheme of single LA domain constructs in the context of the VLDLRΔLBD backbone. (**G**) Raw BLI sensorgrams of biosensors coated with indicated Fc-fusion proteins following incubation with EEEV PE–6 VLPs (*left panel*) or biosensors coated with EEEV PE–6 VLPs following incubation with Fc-fusion proteins in solution (*right panel*). (**I**) Scheme of VLDLR with Trp (W) to Ala (A) mutations in LA1, LA2, LA3, LA5, and LA6 (left) VLDLR (WA), and also VLDLR (WA) with LA4 (F171W), LA7 (R295W), LA8 (K336W) residues changed to Trp. (**J**) SINV-EEEV-GFP PE–6 infection of K562 cells transduced with variants of VLDLR (WA) in which the indicated single LA domain has been reverted to Trp as indicated. (**K**) SINV-EEEV PE–6 GFP infection of K562 cells transduced with variants of VLDLR (WA) in which the indicated LA domains have been reverted to Trp as indicated. (**L**) SINV-EEEV-GFP PE–6 infection of K562 cells transduced with the indicated tandem LA domain constructs in the context of the VLDLRΔLBD backbone. Data in (**B, D, F, I-L**) are pooled from two to six independent experiments. Data in **G** are representative of two independent experiments. *p < 0.05, ****p < 0.0001, n.s., not significant; one-way ANOVA with Dunnett’s post-test.

We next tested N-terminally truncated LA domain variants of VLDLR on K562 cells for their ability to support SINV-EEEV PE–6 infection. After confirming equivalent cell surface expression of the different proteins (**Fig 2C and S3F**), we observed that LA(2–8) or LA(3–8) supported SINV-EEEV PE–6 infection at levels similar to parental VLDLR, whereas partial reductions of infection were seen with cells expressing LA(4–8) or LA(5–8) (**Fig 2D**). However, K562 cells expressing LA(6–8) showed substantially reduced SINV–EEEV infections, and those expressing LA(7–8), LA8, and ΔLBD failed to support infection (**Fig 2D**); thus, LA7 and/or LA8 alone cannot support infection in this context. To identify the LA domains that are functionally relevant for EEEV infectivity, we expressed individual LA domains in the context of the VLDLR ΔLBD backbone with an N-terminal Flag-tag (**Fig 2E and S3G**). After transducing cells, we sorted for similar levels of Flag-expression of all constructs relative to parental VLDLR (**Fig S3H**) and then inoculated them with SINV-EEEV. Expression of LA1, LA2, LA3, LA5, or LA6 on the cell surface promoted infection, albeit much less efficiently than parental VLDLR, whereas LA4, LA7, and LA8 did not support SINV-EEEV PE–6 infection above background (**Fig 2D and F**). These results demonstrate that no single LA domain can support infection as efficiently as the parental 8 LA-domain containing protein, suggesting a requirement for use of multiple LA domains as observed in our structural data.

To corroborate these experiments, we generated single LA-domain Fc fusion proteins and evaluated them for binding to EEEV VLPs using biolayer interferometry (BLI) (**Fig S4A**). We first validated the specificity of the binding assay by immobilizing VLDLR-LBD-Fc (**Fig S4B**) or LDLRAD3-LA1-Fc^40^ on anti-human Fc biosensors. As expected, pins coated with VLDLR-LBD-Fc bound to EEEV, but not VEEV VLPs, and reciprocally, pins coated with LDLRAD3-LA1-Fc bound to VEEV, but not EEEV VLPs (**Fig S4B**). We then tested individual VLDLR LA domains for binding to EEEV. Notably, immobilized VLDLR-LA1, VLDLR-LA2, VLDLR-LA3, VLDLR-LA5, and VLDLR-LA6 captured EEEV VLPs (**Fig 2G**, *left panel*), consistent with the single domain infection experiments. However, EEEV VLPs were unable to bind VLDLR-LA4 or VLDLR-LA7, again consistent with the infection data. To determine how efficiently the single LA domains could bind EEEV VLPs, we immobilized VLPs on mAb-coated biosensors and then evaluated binding of Fc-fusion proteins in solution. In this context, the VLDLR LA domains must bind without the advantage of avidity interactions. In contrast to full-length VLDLR LBD, even at 1 µM concentrations, the single LA domains showed marginal binding to EEEV VLPs in solution (**Fig 2G**, *right panel*).

### Defining the domain binding requirements for efficient EEEV infection

Our data show that EEEV can recognize LA1, LA2, LA3, LA5, and LA6 but not LA4, LA7, or LA8. Sequence alignment of the VLDLR LA domains reveals that the domains recognized by EEEV all share a conserved Trp at the calcium-coordination site, which contrasts with those not recognized by EEEV (LA4, Phe; LA7, Arg; and LA8, Lys), suggesting that a Trp at this position is required for binding (**Fig S3G**). Indeed, prior work showed the Trp residue is critical for LA domain binding of VLDLR to SFV,^46^ and the orientation of the LA domains in our structure suggests an analogous mode of engagement (**Fig 1C**) with density attributable to Trp observed for the LA domain residing within the E1/E2 cleft (interfacing with K156 of EEEV E2) (**Fig 1C**, *inset*). To test this hypothesis, we generated K562 cells expressing a construct in which the Trp of LA1 (W50), LA2 (W89), LA3 (W132), LA5 (W210), and LA6 (W256) of full-length VLDLR were mutated (VLDLR (WA)), (**Fig 2H**). Despite being expressed on the cell surface, VLDLR (WA) was unable to support EEEV infection (**Fig 2I and S4C**), which confirms the functional importance of the Trp residues in these domains. Reciprocally, we mutated the corresponding residues of LA4, LA7, and LA8 to Trp in the context of VLDLR(WA). However, these proteins did not support EEEV infection (**Fig 2I**), suggesting that other LA domain residues contribute to binding, and that Trp substitution alone is not sufficient to promote infection. To demonstrate that VLDLR(WA) could functionally support infection by an alphavirus, we replaced the LA4 domain with LDLRAD3 LA1, which enabled productive SINV-VEEV infection (**Fig S4D**).

Using the full-length VLDLR (WA) as a backbone, we next generated constructs in which Trp was re-introduced into each LA domain to assess function. We observed low but detectable levels of EEEV infection in cells expressing constructs in which only one LA domain encoded the conserved Trp (**Fig 2J and S4E**), which was consistent with our single-LA domain binding experiments (**Fig 2F**). Given our structural data (**Fig 1C**), we next tested whether the presence of two functional LA domains (re-inserted Trp residues) in the context of full-length VLDLR (WA) would enhance infection. While some (*e.g.*, LA2 + LA5, LA2 + LA6, and LA5 + LA6) constructs supported lower levels of infection, others (*e.g.,* LA1 + LA2, LA1 + LA3, and LA3 + LA5) mediated infection at levels similar to the parental VLDLR (**Fig 2K and S3F**). The connected LA densities in our structure suggested that EEEV may engage two contiguous LA domains simultaneously. To probe this possibility, we expressed “mini-receptors” harboring tandem LA domains [LA(1–2), LA(2–3), LA(3–4), LA(4–5), LA (5–6), and LA(6–7)] on the VLDLRΔLBD backbone (**Fig 2L and S4G**). Notably, LA(1–2) ΔLBD supported similar levels of EEEV infection compared to parental VLDLR (**Fig 2L**), much like the full-length VLDLR (WA) with LA1 and LA2 intact (**Fig 2K**). Lower levels of infection were observed with the other tandem domain “mini-receptors” with LA(2–3)ΔLBD barely supporting EEEV (**Fig 2L**). We confirmed LA(2–3)ΔLBD was functional as it supported SINV-SFV infection (**Fig S4H**). Coupled with our binding data (**Fig 2G**), these results suggest that while LA(2–3) may support attachment of EEEV, it inefficiently promotes viral entry. Overall, these experiments with two LA domain constructs support the idea that concurrent interactions with multiple LA domains are required to support efficient EEEV infection.

### LA domain decoy receptors can neutralize EEEV infection in vitro

As another measure of how EEEV engages multiple LA domains, we performed neutralization assays with soluble decoy receptors comprised of different VLDLR LA domains. To assess neutralization, Fc-fusion proteins (10 μg/mL) were incubated with SINV-EEEV PE–6 prior to inoculation of 293T cells. Whereas VLDLR-LBD-Fc neutralized SINV-EEEV PE–6 almost completely, the single LA domain-Fc proteins did not (**Fig 3A**), which correlated with their relatively poor binding in solution by BLI (**Fig 2G**). We then performed neutralization with different N-terminal fragments of the VLDLR receptor (LA(1–2), LA(1–3), LA(1–4), LA(1–5), and LA(1–6)) fused to an Fc domain. Notably, all N-terminal fragments neutralized infection, with increased potency observed with longer fragments that contain more LA domains (**Fig 3B**). The results with LA(1– 2)-Fc are consistent with the sufficiency of these two VLDLR LA domains to support efficient SINV-EEEV infection, although the potency of neutralization was greater for decoy receptor proteins expressing additional LA domains that possibly could engage all three binding sites. We extended this analysis by testing additional two domain constructs (**Fig 3C**). Whereas LA(1–2)- Fc and LA(2–3)-Fc completely neutralized infection at 10 μg/mL, LA(3–4)-Fc, LA(4–5)-Fc, and LA(5–6)-Fc did not. We also tested C-terminal fragments of VLDLR (LA(2–8)-Fc, LA(3–8)-Fc, LA(4-6)-Fc) for their ability to neutralize SINV-EEEV PE–6 infection. As LA(2-8)-Fc, LA(3-8)- Fc, and LA(3-6) efficiently inhibited SINV-EEEV PE–6 infection (**Fig 3D**), binding and neutralization of EEEV PE–6 virions in solution does not require LA1 or LA2, consistent with efficient infection mediated by combinations of other domains (*e.g.*, LA3 + LA5) (**Fig 2K**).

**Figure 3.**
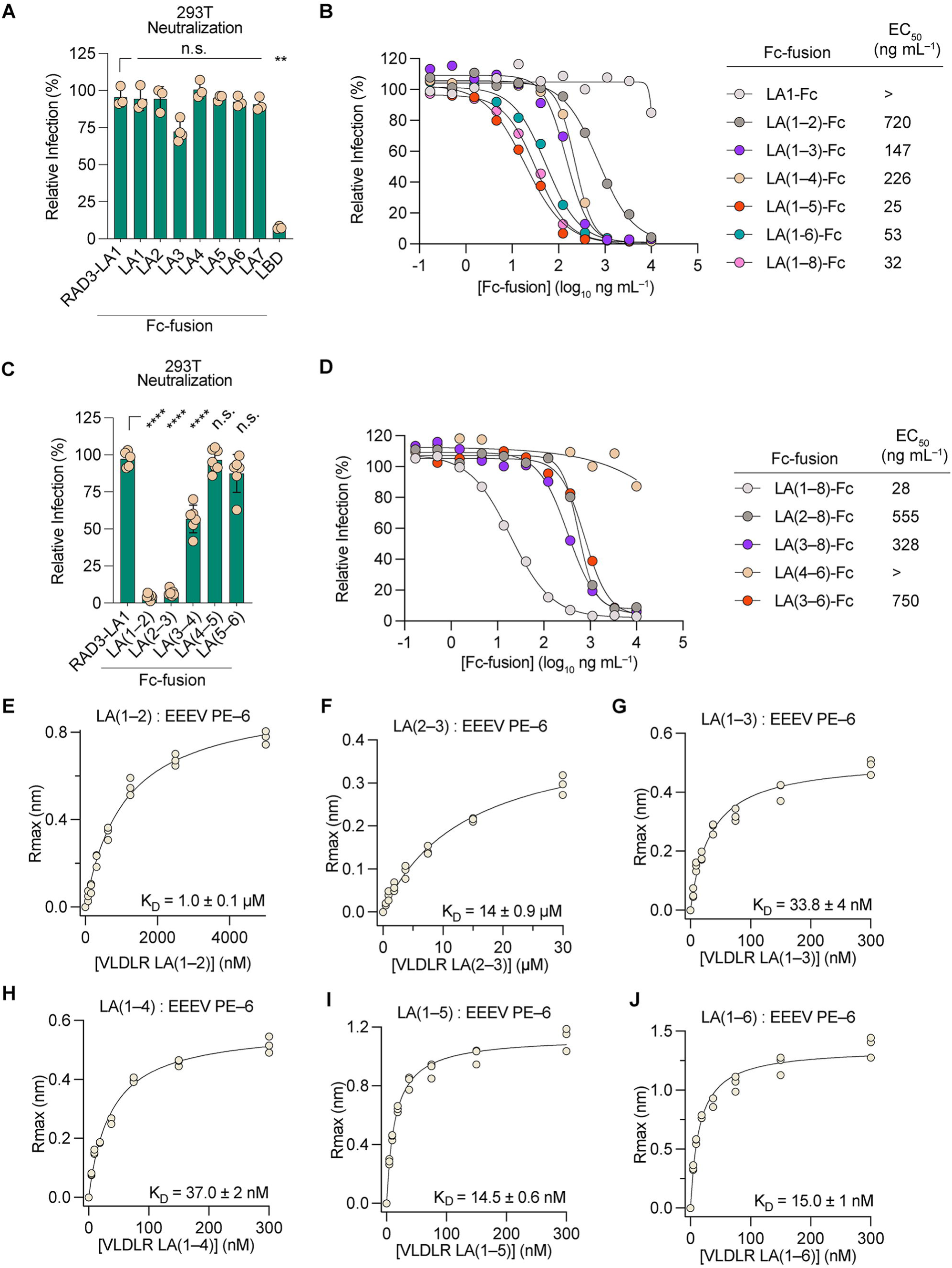
Multiple LA domains mediate neutralization by VLDLR receptor decoys against EEEV. (**A, C**) Infection of 293T cells by SINV-EEEV-GFP PE–6 following pre-incubation with the indicated Fc-fusion proteins (10 μg/mL) prior to inoculation. GFP expression was measured by flow cytometry. (**B, D**) Dose-response curves of neutralization by the indicated Fc-fusion proteins against SINV-EEEV-GFP PE–6. Data in (**A, C**) are pooled from three to six independent experiments performed in technical duplicate. **p < 0.01, **** p < 0.0001, n.s., not significant by one-way ANOVA with Dunnett’s post-test. Data in (**B, D**) is representative of three independent experiments with mean half-maximal effective inhibitory concentrations (EC_50_ values) calculated. (**E–J**) Steady-state BLI curves of the indicated monovalent VLDLR domains bound to EEEV PE–6 VLP coated biosensors. Data are pooled from three independent experiments.

Our results showed that Fc decoy molecules expressing tandem but not single LA domains were sufficient to neutralize SINV-EEEV infection. To better understand the functional hierarchy of LA domain engagement, we performed steady-state affinity analyses via BLI with monovalent forms of LA(1–2) and LA(2–3), as these molecules neutralized SINV-EEEV infection when expressed as Fc fusion proteins. Whereas LA(2–3) bound EEEV PE–6 VLPs with low affinity (K_D_, 14 ± 0.9 μM), LA(1–2) bound with 10-fold higher affinity (K_D_, 1.0 ± 0.1 μM) (**Fig 3E-F**). We then evaluated the binding of LA(1–3), LA(1–4), LA(1–5), and LA(1–6) to determine whether the addition of more C-terminal LA domains would enhance affinity (**Fig 3G-J**). Indeed, addition of LA3 dramatically improved binding, with LA(1–3) and LA(1–4) exhibiting K_D_ values of 33.8 ± 4 and 37.0 ± 2 nM, respectively. The observed enhancement of affinity plateaued with the addition of LA5 and LA6, as LA(1–5) and LA(1–6) bound with respective K_D_ values of 14.5 ± 0.6 nM and 15.0 ± 1 nM. Although it is possible that these affinities reflect some receptor fragments spanning immobilized VLPs (rather than simply binding multiple sites on a single VLP), the data are consistent with our structural, neutralization, and infection studies, further suggesting that EEEV can engage multiple LA domains concurrently with higher avidity. Taken together, our tandem LA domain neutralization (**Fig 3C**), affinity measurements (**Fig 3E-J**), and “mini-receptor” infection (**Fig 2L**) studies indicate that while EEEV PE–6 can redundantly bind other LA domains, it engages VLDLR LA(1–2) domains particularly well.

### Cryo-EM structures of EEEV VLP in complex with VLDLR fragments

To understand how EEEV can engage different LA domains of VLDLR, we attempted to reconstruct EEEV VLPs in complex with different VLDLR fragments using cryo-EM. We first assessed the structures of tandem sequential LA domains bound to EEEV VLPs, as we observed a clear linking density between the cleft and A domain site in our full-length VLDLR-EEEV structure. We focused on LA(1–2) and LA(2–3), as these proteins (as Fc-fusions) neutralized SINV-EEEV PE–6 infection, and also included LA(5–6) as both individual domains can interact with EEEV (**Fig 2F-G**). For uncertain reasons, we observed no receptor density for LA(2–3) or LA(5–6), as either monomeric proteins or bivalent Fc fusions, despite incubation of EEEV VLPs with ten-fold molar excess of receptor in solution. However, when EEEV VLPs were incubated with VLDLR LA(1–2), two LA domains with a linking density were observed with high occupancy (**Fig 4A**), matching the connected domains observed in the full-length VLDLR structure (**Fig 1C**). To better define this interaction, we performed focused refinement of the asymmetric unit to 3.09-Å resolution (**Fig 4A**, **S1, and S2C**), enabling delineation of most sidechains at the binding interface (**Fig S2D**). VLDLR LA1 binds within the cleft between E1/E2 heterodimers, establishing conventional “wrapped” and “intraspike” contacts to neighboring heterodimers (**Fig 4B-C**),^41,43^ whereas LA2 assumes an angled position atop the A domain of EEEV E2, a site not occupied in other alphavirus-receptor structures (**Fig 4C**). At the wrapped interface, VLDLR LA1 engages residues of the conserved fusion loop of EEEV E1 (residues 85, 87-92, 95, 97, and 98) (**Fig 4D**). On the opposing face of the cleft, LA1 forms electrostatic interactions with residues H155-K156-R157 of the intraspike E2 (**Fig 4E**). This electropositive “HKR” loop of EEEV E2 targets the highly conserved calcium-coordination site of LA1, establishing salt-bridges with VLDLR residues D53, D55, and D57, as well as hydrophobic and cation-π interactions with W50 (**Fig 4E**). EEEV also targets the calcium cage of VLDLR LA2, coordinating via E2 residue K231 with VLDLR residues D92, D94, and D96, as well as W89; LA2 is also stabilized by K232 and an additional loop (residues 56-59) from the A domain of E2 (**Fig 4F**). Cryo-EM processing statistics and a complete list of interfacial residues for EEEV and VLDLR LA(1–2) are provided (**Tables S1 and S2**).

**Figure 4.**
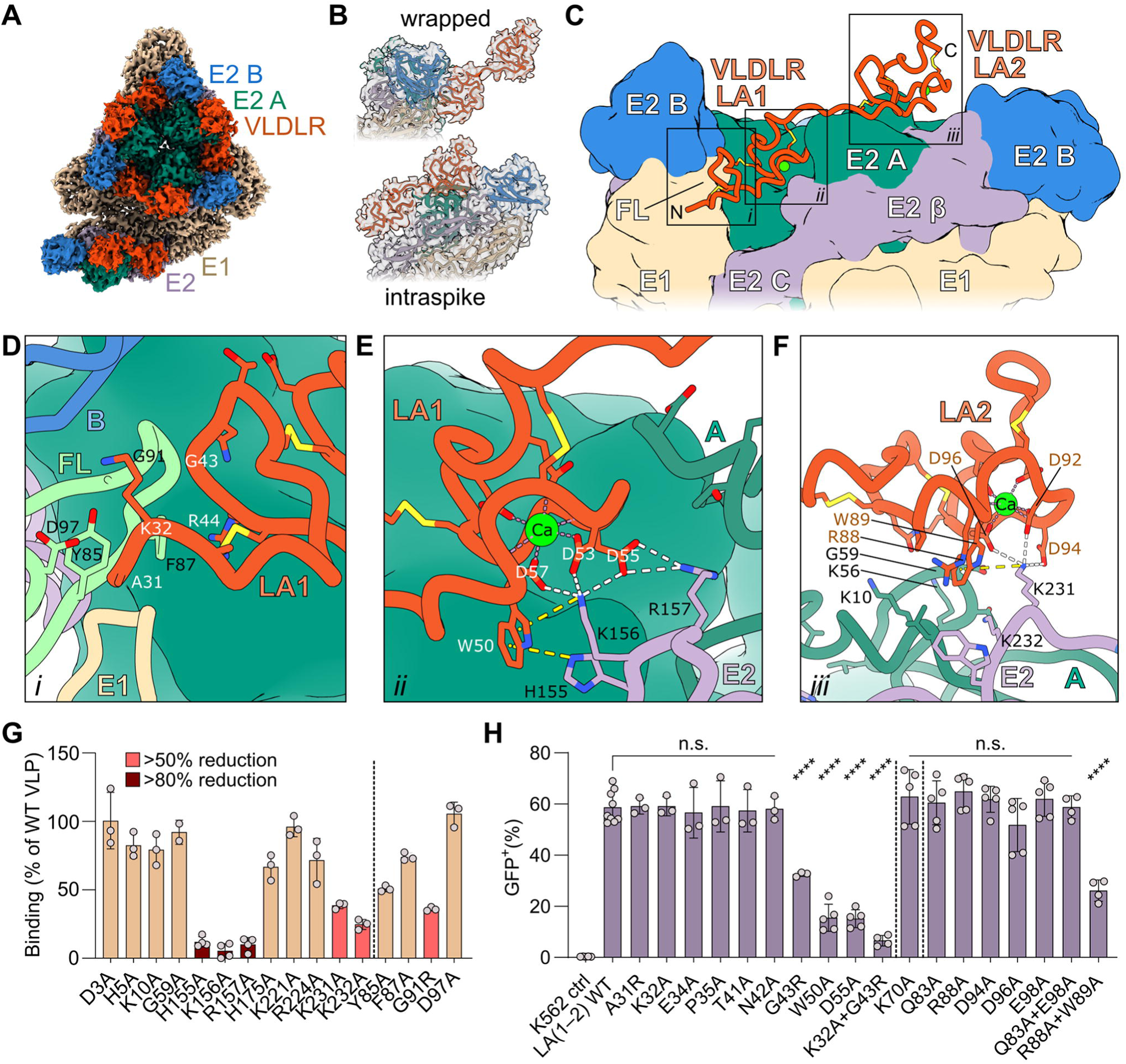
Cryo-EM structure of EEEV PE–6 VLPs in complex with VLDLR LA(1– 2). (**A**) Focused reconstruction of the EEEV PE–6 asymmetric unit in complex with VLDLR-LA(1–2). Proteins are differentially colored with E1 in tan, E2 A domain in sea green, E2 B domain in blue, the remainder of E2 colored purple, and VLDLR shown in orange. (**B**) Individual E1/E2 heterodimers at the binding interface, illustrating conventional “wrapped” and “intraspike” contacts, colored as in (**A**).^41,43^ (**C**) A ribbon diagram of VLDLR LA(1–2) (orange) overlaying a surface representation of neighboring E1/E2 heterodimers, colored as in (**A**). Domains of E2 are labeled, as is the fusion loop (FL) of E1. (**D-F**) Magnified regions from boxes in panel (**C**). Interface details between VLDLR LA1 and the E1 fusion loop (pale green) (**D**), LA1 and E2 (**E**), and LA2 and E2 (**F**) are shown. VLDLR residues are labeled in white or orange, with EEEV residues labeled in black. Predicted salt bridges (interatomic distance ≤ 4.0 Å) and cation-π interactions (≤ 6.0 Å from aromatic plane) are demarcated by white or yellow dashed lines, respectively. (**G**) Binding of LA(1–2)-Fc to captured wild-type (WT) and mutant EEEV FL93–939 VLPs. Biosensors were coated with WT or mutant VLPs followed by incubation with 1 μM of LA(1–2)-Fc for 300 sec. Binding was calculated as percent signal (R_max_) relative to WT VLPs. (**H**) Infection of K562 cells expressing WT and mutant constructs of VLDLR LA(1–2) by SINV-EEEV PE–6. Infection was assessed by GFP expression using flow cytometry. Data in **G-H** are pooled from two to four independent experiments. **p < 0.01, ****p < 0.0001, n.s., not significant by one-way ANOVA with Dunnett’s post-test.

To corroborate our structural model of EEEV in complex with LA(1–2), we selected clones from a previously generated VLP expression library^50^ (EEEV strain FL93–939) encoding alanine mutations in the E2 gene and measured binding to VLDLR LA(1–2)-Fc protein by BLI. Compared to wild-type EEEV VLPs, an almost complete loss of binding of VLDLR LA(1–2)-Fc was observed with EEEV VLPs encoding E2-H155A, E2-K156A, or E2-R157A substitutions, and a partial loss of binding (>50% reduction) was measured with E2-K231A or E2-K232A mutants (**Fig 4G**). Several other E2 substitutions (D3A, H5A, K10A, G59A, H175A, K221A, R224A) did not impact EEEV VLP binding, suggesting a less critical role for these residues **(Fig 4G**). To assess the interaction of the LA1 domain with the fusion loop in E1, we generated additional mutant VLPs, including E1-Y85A, E1-F87A, E1-G91R, and E1-D97A, and evaluated binding of LA(1–2)-Fc; we observed >50% reduction in binding for E1-G91R (**Fig 4G**), likely due to steric hindrance at this site or reduced dihedral flexibility of the fusion loop.

To further evaluate our model, we generated structure-guided mutants in both LA1 and LA2 domains in the context of the LA(1–2)ΔLBD “mini-receptor” and assessed their effects on SINV-EEEV PE–6 infection. Of the 9 mutants we made in the LA1 domain, only G43R, W50A, and D55A substitutions resulted in decreased SINV-EEEV PE–6 infection (**Fig 4H and S4I**). A K32A/G43R double substitution showed greater reduction in infectivity, consistent with the LA1–E1 fusion loop interaction in our model and the impaired binding of LA(1–2)-Fc to E1-G91R (**Fig 4D and H**). Although the single LA2 domain substitutions we tested (K70A, Q83A, R88A, or D94A) did not impair the ability of LA(1–2)ΔLBD to support infection, loss of infectivity was observed with R88A + W89A in LA2, again confirming the importance of the conserved Trp (**Fig 4H**).

Given that we were unable to visualize LA(2–3) or LA(5–6) via cryo-EM, we characterized how EEEV engages LA3, LA5, and LA6 by complexing EEEV PE–6 VLPs with LA(1–3), LA(1–5mut3) (harboring a W132A mutation in LA3), or LA(1–6mut3,5) (harboring W132A and W210A mutations in LA3 and LA5, respectively), as LA(1–2) could anchor VLP binding to the E1/E2 cleft and E2 A domain sites (**Table S1**). Each of these reconstructions showed a density, albeit not fully occupied, at the lateral E2 B domain site (**Fig 5A**), suggesting that LA3, LA5, or LA6 can engage this site interchangeably. The more highly occupied density observed at the B domain site for full-length VLDLR might reflect the collective interactions of multiple domains (*e.g*., LA3, LA5, or LA6) (**Fig 1A-B**). To verify that the B domain density in our full-length VLDLR structure can be attributed to LA3, LA5, and LA6, we reconstructed EEEV VLPs in complex with VLDLR LBDmut3,5,6 (harboring W132A, W210A, and W256A in LA3, LA5, and LA6, respectively). As expected, we observed only LA1 and LA2 domains with no visible density at the lateral B domain site (**Fig 5A**).

**Figure 5.**
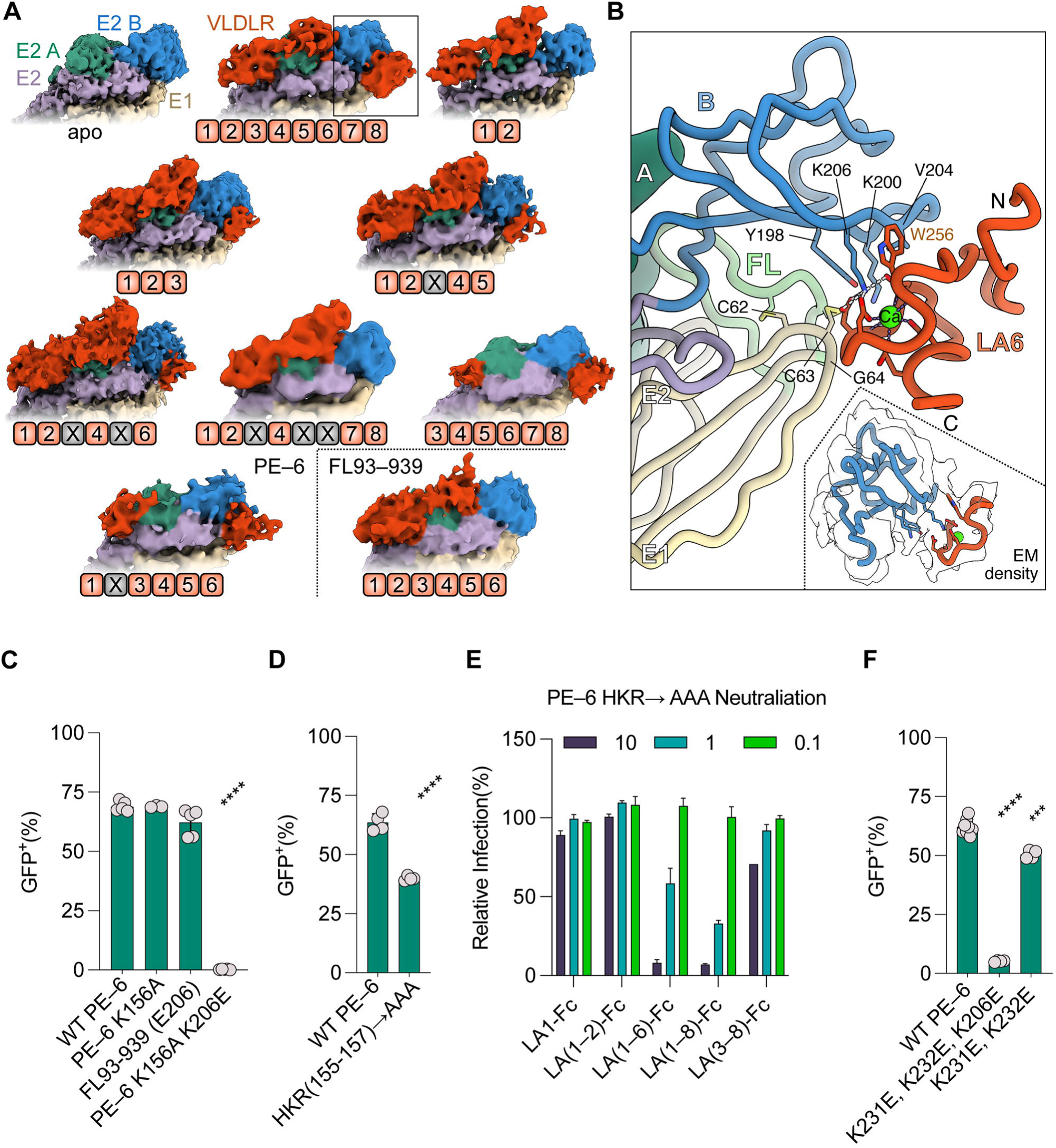
Mapping LA domain binding and EEEV E2 binding sites. (**A**) Cryo-EM reconstructions of EEEV PE–6 or FL93–939 VLP in complex with different VLDLR fragments, identified schematically below each reconstruction. VLDLR constructs include full-length VLDLR; LA(1–2); LA(1–3); LA(1–5mut3); LA(1–6mut3,5); LBDmut3,5,6; LA(3–8); LA(1–6mut2); and LA(1–6). EEEV E1 is shown in tan, E2 A domain in sea green, E2 B domain in blue, the remainder of E2 in purple, and VLDLR in orange. (**B**) A ribbon diagram depicting the interaction of VLDLR LA6 with EEEV E1 and E2, colored as in (**A**) with the fusion loop (FL) shown in pale green. Predicted salt bridges (interatomic distance ≤ 4.0 Å) are demarcated by white dashed lines. The fit of this model within the experiment density is pictured within the inset. (**C, D, F**) Infection of SINV-EEEV-GFP PE–6 WT and mutants in K562 cells expressing WT VLDLR as quantified by GFP expression using flow cytometry. (**E**) Dose-response neutralization (10, 1, and 0.1 μg/mL) of indicated Fc-fusion proteins against SINV-EEEV PE–6 HKR → AAA virus in 293T cells. Infection was assessed by GFP expression using flow cytometry. Data in (**C, D, F**) are pooled from three to six independent experiments. Data in **E** are representative are pooled from two independent experiments. ***p < 0.001, ****p < 0.0001; one-way ANOVA with Dunnett’s post-test.

Since no single LA domain engaged the E2 B domain site with full occupancy, we used the B domain density produced by full-length VLDLR to build LA6 (as an example) for our interface analysis. In our model, LA6 principally engages E2-K206 via W256, D259, D261, and D263 within the calcium-coordination site (**Fig 5B**). VLDLR may contact other residues of E2 (*e.g.*, Y198, K200, and V204) or of E1 domain II (*e.g.*, C62, C63, and G64) at this site; however, at a moderate contour threshold, only the calcium-coordination site of the LA domain is visible in our map (**Fig 5B** and **S2D**). The minimal nature of this interface (< 300 Å^2^ buried surface area, with ∼120 Å^2^ solely on K206) may explain the LA domain promiscuity observed at this binding site as well as the relative difficulty of resolving a single domain at this position.

To further define the patterns of VLDLR domain binding to EEEV, we generated cryo-EM reconstructions of EEEV VLPs in complex with LA(3–8) or LA(1-6mut2) (harboring W89A in LA2) (**Table S1**). For both reconstructions, we observed densities in the cleft between E1-E2 heterodimers and on the side of E2 B domain, whereas density was absent in the E2 A domain (**Fig 5A**). This result indicates the glycoprotein cleft of EEEV can be recognized by different LA domains (including non-LA1 domains), whereas the E2 A domain appears to exhibit relative specificity for LA2, as other domains are not visualized at this site. Taken together, our structural analysis shows that multiple sites on the EEEV glycoprotein (E2/E1 cleft, E2 A domain, E2 B domain) can recognize different VLDLR LA domains with some degree of promiscuity. Notwithstanding this point, our results suggest that EEEV engages LA1 and LA2 efficiently at the cleft and A domain sites, as these are the only LA domains fully and unequivocally visualized by cryo-EM with either full-length or fragments of VLDLR; indeed, we did not observe other LA domains occupying the cleft or A domain sites in the absence of LA(1–2). The relative importance of LA(1–2) is consistent with our functional studies, as they are the only tandem LA domains that support efficient SINV-EEEV infection, neutralize EEEV infection as Fc fusions, and bind with moderate affinity as monovalent proteins.

### EEEV strains do not conserve all VLDLR-binding sites

Given that cell surface expressed LA(1–2) protein efficiently supports SINV-EEEV infection and LA(1–2)-Fc potently neutralizes SINV-EEEV infection, we considered whether there was redundancy in the three binding sites (E1/E2 cleft, E2 A domain, and E2 B domain), and that any given site might not be required to mediate VLDLR-dependent interactions. We first evaluated whether the basic amino acids interacting with Trp of the LA domains were conserved in EEEV strains. The E2-HKR loop (H155-K156-R157; cleft) and E2-K/R231 + E2-K232 (A domain shelf) residues are conserved in all 692 EEEV structural polyprotein sequences present in GenBank (**Table S3**). However, the E2 B domain binding site, which features residue E2-K206 in the PE–6 strain, is not conserved in FL93–939 (E206), FL91–469 (E206), or 690 other EEEV strains; residue E2- K206 is present only in one other EEEV strain (A61–1K) and in one EEE complex Madariaga virus strain (BeAr348998) (**Table S3**). To confirm that the E2-E206 substitution abrogates VLDLR engagement at the B domain site, we performed cryo-EM with VLDLR LA(1–6) complexed with EEEV FL93–939 VLPs; this strain differs from PE–6 in E2 only at the E2-E206 residue. As expected, we did not observe receptor density at the B domain site for FL93–939 (**Fig 5A**). Moreover, VLDLR LA(1–2)-Fc neutralized SINV-EEEV FL93–939 infection equivalently compared to LA(1–3)-Fc, LA(1–4)-Fc, LA(1–5)-Fc, LA(1–6)-Fc, or LBD-Fc (**Fig S5A**), providing supporting evidence that LA3, LA5, and LA6 do not engage a third site on FL93–939. Given that a prior study showed VLDLR-dependent infection with reporter virus particles encoding FL91–469 structural genes,^45^ the E2-K206 residue in EEEV PE–6 may be dispensable or act as a gain-of-function. As K562 cells ectopically expressing VLDLR are permissive to SINV-EEEV FL93–939 (**Fig 5C**), engagement of the E2 B domain site on EEEV is not required for infectivity.

The E2-206K residue might provide an advantage for EEEV PE–6 by enhancing its binding for VLDLR relative to EEEV FL93–939. To test this possibility, we performed affinity measurements of LA(1–6) in complex with FL93–939 or FL93–939 engineered with an E2- E206K substitution. We observed a slightly lower affinity of LA(1–6) binding for FL93–939 (K_D_, 50.0 ± 4.7 nM) compared to PE–6 (15.0 ± 1 nM), but this difference was abrogated by the E2-E206K change in FL93–939 (K_D_, 17.4 ± 2.5 nM). These results suggest that EEEV PE–6 has a small but measurable advantage in VLDLR-binding that is due to the E2-206K polymorphism (**Fig S5B**).

Beyond binding affinity, the E2-E206K substitution might enable more versatile domain usage, as other LA domains (*e.g.*, LA3, LA5, and LA6) could be engaged through the B domain binding site. To begin to address this possibility, we evaluated whether LA3, LA5, and/or LA6 could bind and support infection of EEEV FL93–939. When immobilized on a biosensor in the solid phase, the single LA domain-Fc proteins could capture FL93–939 VLPs (**Fig S5C**), and K562 cells transduced with VLDLR (WA) constructs with intact LA3, LA5, or LA6 domains supported low levels of SINV-EEEV FL93–939 infection (**Fig S5D**). When VLDLR(WA) constructs were expressed on the cell surface with two intact LA domains (*e.g.,* LA1+LA3, LA1+LA5, or LA1+LA6), more efficient SINV-EEEV FL93–939 infection was observed, comparable to that seen with LA1+LA2 (**Fig S5E**); however, and in contrast to SINV-EEEV PE–6, domain combinations without LA1 did not support robust SINV-EEEV FL93–939 infection. The greater dependence of FL93–939 on VLDLR LA1 suggests that while non-LA1 domains can enter the E1/E2 cleft (as observed in the LA(3–8) reconstruction (**Fig 5A**)), they engage this site poorly and cannot support efficient infection without assistance of the B domain VLDLR-binding site. We extended this analysis by generating mutant LA(1–6)-Fc decoys with Trp mutations to assess binding as a function of neutralization activity. Consistent with our infection data, a W50A substitution in LA1 (LA(1–6mut1)) impaired the neutralizing activity of LA(1–6)-Fc against SINV-EEEV FL93–939 but not against SINV-EEEV PE–6 (**Fig S5F-G**). However, a W89A substitution in LA2 (LA(1–6mut2)) did not lessen the neutralizing activity of LA(1–6)-Fc against either SINV-EEEV FL93–939 or PE–6. To determine which other LA domains contributed to neutralization of FL93–939, we generated LA(1–6mut2) with additional Trp mutations alone or in combination with the other LA domains. As expected, LA(1– 6mut2,3,5,6) did not neutralize SINV-EEEV FL93–939 infection (**Fig S5H**). In contrast, LA(1– 6mut2,5,6) and LA(1–6mut2,3,5) neutralized SINV-EEEV FL93–939 infection, albeit less potently than LA(1–6mut2), suggesting that LA3 and LA6 also can engage the A domain site.

To corroborate that a non-LA2 domain can recognize the A domain site, we performed semi-quantitative BLI to evaluate the binding of LA(1–6mut2)-Fc to EEEV FL93–939 VLPs with or without K231E and K232E substitutions in E2. As LA(1–6mut2) showed substantially less binding to FL93–939 E2-K231E/E2-K232E VLPs than WT FL93–939 VLPs, non-LA2 domains of VLDLR likely can bind the A domain site of EEEV FL93–939 (**Fig S5I**).

### Two sites on EEEV E2 are required for efficient VLDLR-dependent infection

To test the functional importance of the cleft binding site, we generated a SINV-EEEV PE–6 E2-K156A virus. This mutant virus infected VLDLR-expressing K562 cells as efficiently as the parental SINV-EEEV PE–6 (**Fig 5C**). We then generated a SINV-EEEV PE–6 E2-K156A/E2-K206E double mutant virus in which both the cleft and B domain binding sites are inactivated. While this mutant virus grew as efficiently as parental SINV-EEEV PE–6 in BHK-21 and Vero cells, the SINV-EEEV-PE–6 E2-K156A/E2-K206E mutant virus did not infect VLDLR-expressing K562 cells (**Fig 5C**). Thus, the A domain site (E2-K231 and E2-K232) alone is insufficient to mediate VLDLR-dependent infection. These data suggest that E2-K206 on the residue B domain site can rescue VLDLR-dependent infection in the event of mutations within the cleft.

We next considered whether the neighboring residues E2-H155 and E2-R157 in the cleft could compensate for the E2-K156A mutation in the context of the SINV-EEEV PE–6 strain, given that SINV-EEEV PE–6 E2-K156A used VLDLR as efficiently as the parental virus (**Fig 4H**). To test whether cleft binding was incompletely abolished with the E2-K156A mutant, we generated a SINV-EEEV PE–6 HKR→AAA virus. This mutant virus showed only a moderate loss-of-infection phenotype compared to parental SINV-EEEV PE–6 in VLDLR-expressing K562 cells (**Fig 5D**), suggesting that the cleft binding site is not essential for EEEV PE–6 to utilize VLDLR as a receptor. Moreover, when we performed neutralization experiments with SINV-EEEV PE–6 HKR→AAA virus with Fc fusion protein decoys, we observed inhibition with VLDLR-LBD-Fc (**Fig 5E**), which in theory can still recognize two binding sites (E2 A and B domains) on this mutant virus. In contrast, LA(1–2)-Fc or LA(3–8)-Fc failed to neutralize SINV-EEEV PE–6 HKR→AAA (**Fig 5E**); in our cryo-EM reconstructions, LA(1–2) or LA(3–8) bind only one other site in addition to the E1/E2 cleft (E2 A and E2 B domains, respectively). Together, these results indicate the E1/E2 cleft site is not absolutely required to efficiently bind VLDLR for EEEV strains featuring E2-K206.

We also tested whether the cleft binding site alone was sufficient for VLDLR-dependent infection by generating SINV-EEEV PE–6 E2-K206E/E2-K231E/E2-K232E, in which the E2 A and B domain binding sites are inactivated. While this mutant virus grew in BHK-21 cells, it poorly infected VLDLR-expressing K562 cells (**Fig 5F**). We extended these studies by generating SINV-EEEV PE–6 E2-K231E/E2-K232E (K206 and cleft binding site present); this mutant virus used VLDLR for infection with only a slightly reduced efficiency (**Fig 5F**). These results are consistent with a requirement for two distinct binding sites on EEEV (cleft + A domain, cleft + B domain, or A + B domains) for efficient VLDLR-dependent infection.

### Comparison of receptor engagement by alphaviruses

The structural interface of EEEV with VLDLR resembles yet differs from those observed for other alphavirus-receptor complexes (**Fig S6-S7**). Whereas LA1 binds within the cleft between E1/E2 heterodimers in a manner analogous to VEEV/LDLRAD3 and CHIKV/MXRA8 (**Fig 6A-C**),^41–44^ LA2 engages a unique site on a shelf atop the A domain of EEEV E2 and other LA domains engage a third site on E2 B domain in some EEEV strains (**Fig 6A**). In comparison, SFV engages VLDLR LA3 outside of the cleft through domain III of E1 (**Fig 6D**).^46^ Like EEEV, SFV can engage multiple domains of VLDLR;^46^ however, the concomitant engagement of different LA domains was not structurally or functionally resolved, and unlike EEEV, SFV uses a single site in E1 for receptor binding.

**Figure 6.**
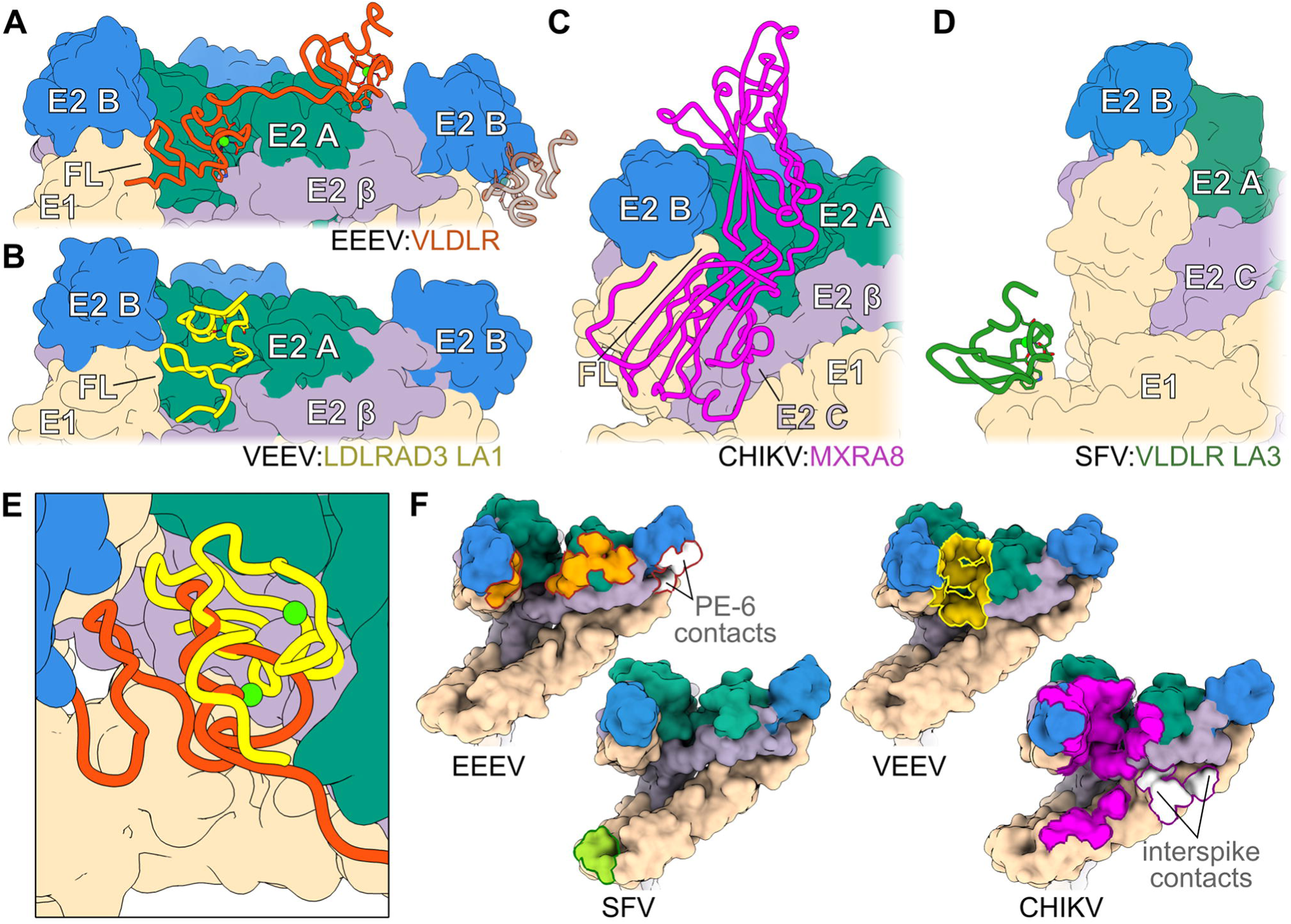
Comparative analysis of virus-receptor complexes. (**A-D**) Structures of known alphavirus-receptor complexes, with VLDLR (orange; strain-specific site shown as transparent), LDLRAD3 LA1 (yellow, PDB 7N1H), MXRA8 (magenta, PDB 6NK7), and VLDLR LA3 (green, PDB 8IHP) displayed as ribbon diagrams, overlaying surface representations of EEEV, VEEV, CHIKV, or SFV structural proteins. E1 is shown in tan, E2 A domain in sea green, E2 B domain in blue, and the remainder of E2 (including C domain and the β-linker) in pale purple. (**E**) Magnified top view showing overlay of VLDLR LA1 (orange) and LDLRAD3 LA1 (yellow) within the EEEV/VEEV receptor binding cleft. (**F**) Surface representations of neighboring E1/E2 heterodimers, colored as in (**A-D**), with receptor binding interfaces (determined by Proteins, Interfaces, Structures, and Assemblies (PISA))^51^ highlighted on their respective alphaviruses.

Although EEEV and VEEV can engage LA domains using their E1/E2 clefts in a roughly similar location, EEEV establishes two limited points of contact with VLDLR LA1 using the fusion loop of E1 and the HKR loop of E2. LA1 is otherwise suspended away from the back of the cleft (**Fig 6E**), burying only ∼650 Å^2^ of the viral surface (by PISA^51^ analysis) (**Fig 6F**). In contrast, VEEV positions LDLRAD3 LA1 inward to form extensive hydrophobic and Van der Waals contacts at the back of the cleft (∼900 Å^2^) (**Fig 6E-F**).^43,44^ Accounting for the two domains together, VLDLR LA1 and LA2 establish an interface larger (∼1,100 Å^2^) than that of LDLRAD3, and an additional ∼290 Å^2^ is buried at the B domain site in some EEEV strains (**Fig 6F**). This three-site interface still is smaller than the expansive interaction between MXRA8 and CHIKV (∼2,200 Å^2^), and the single VLDLR-binding site on SFV is even more limited (∼380 Å^2^) (**Fig 6F**). Nonetheless, because EEEV and SFV can engage multiple LA domains simultaneously,^46^ they bind VLDLR with higher effective affinity (K_D_ values of ∼15 nM and ∼2 nM^46^, respectively) than CHIKV/MXRA8 (K_D_ of 84 to 270 nM) or VEEV/LDLRAD3-LA1 (K_D_ of 50 nM). However, EEEV’s usage of multiple glycoprotein sites for avid receptor engagement differs from SFV and from all previously described alphavirus-receptor complexes.

### LA(1–2)-Fc protects against lethal EEEV challenge in a mouse model

We next tested the protective efficacy of the VLDLR-LBD-Fc decoy and LA(1–2)-Fc against EEEV *in vivo*. We used LA(1–2)-Fc as it retains neutralizing activity against EEEV and lacks other LA domains of VLDLR (*e.g.*, LA5 and LA6) that are implicated in endogenous lipoprotein binding,^52^ which could impact bioavailability and efficacy. We administered PBS or 100 μg (∼5 mg/kg) of LDLRAD3-LA1-Fc, VLDLR LBD-Fc, or VLDLR LA(1–2)-Fc to female CD-1 mice 6 h prior to subcutaneous inoculation with 10^3^ FFU of the authentic CDC Select Agent EEEV FL93–939 strain. All animals treated with PBS, LDLRAD3-LA1-Fc, or LBD-Fc died within 8 days of infection (**Fig 7A**). In contrast, mice given VLDLR LA(1–2)-Fc were completely protected from lethality and did not show weight loss or signs of disease (ruffled fur, hunched posture, seizures, or moribundity) (**Fig 7B-C**). Although LBD-Fc exhibited similar neutralization potency as LA(1–2)-Fc against FL93–939, its lipoprotein binding activity *in vivo* may have limited efficacy. Given the protective efficacy of LA(1–2)-Fc against subcutaneous EEEV infection, we next tested its function in the more stringent aerosol challenge model. Mice administered VLDLR LA(1–2)-Fc were substantially protected (70% vs 0% survival) from lethal aerosol EEEV FL93– 939 infection compared to LDLRAD3-LA1-Fc treated mice (**Fig 7D**). Thus, and in contrast to a full-length receptor decoy, these data highlight the protective activity of a truncated two-domain VLDLR decoy molecule against EEEV infection.

**Figure 7.**
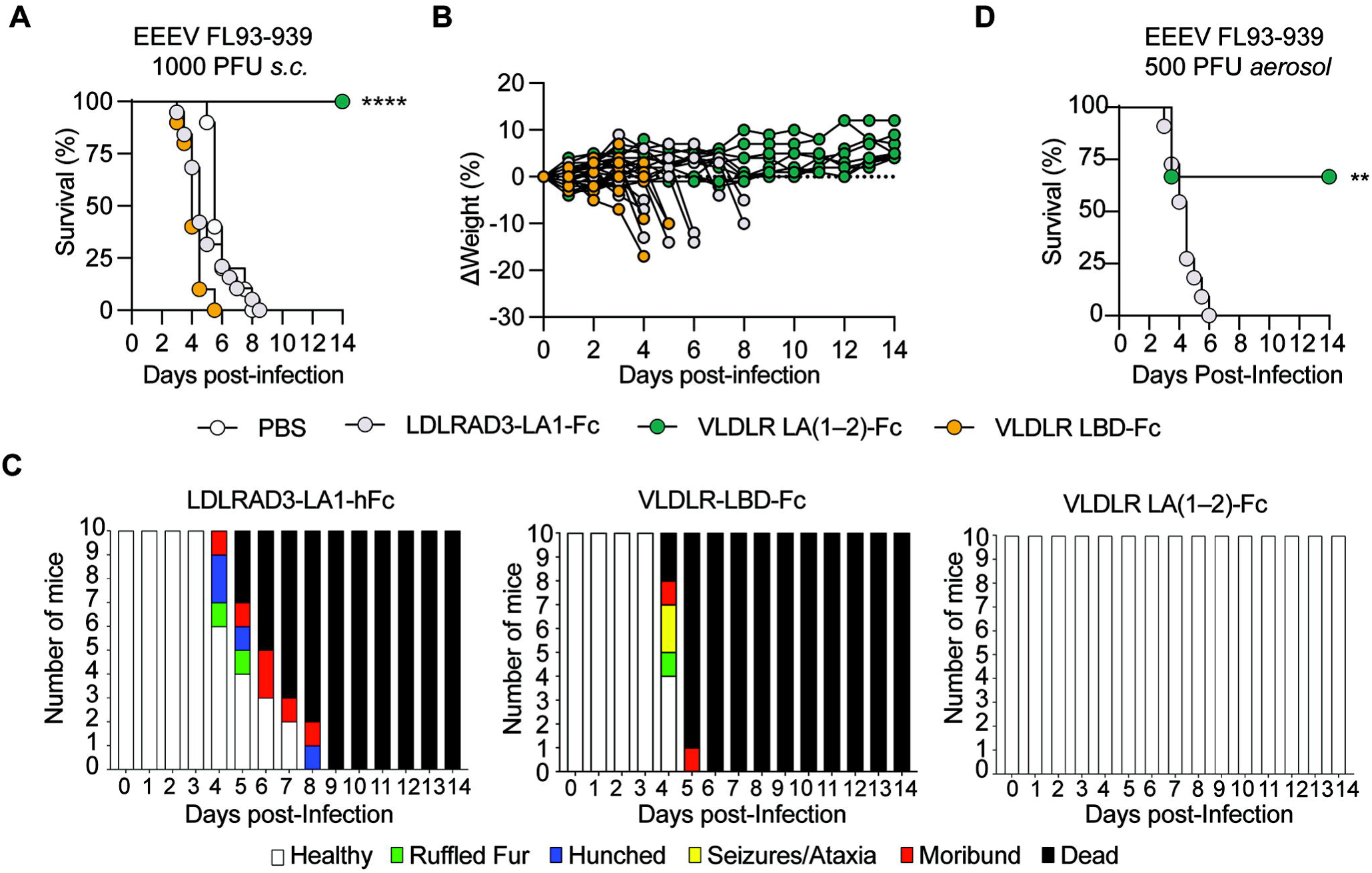
VLDLR LA(1–2)-Fc protects mice against EEEV FL93–939 challenge *in vivo*. (**A–C**) Survival (**A**), weight change (**B**), and clinical scores (**C**) of 5-7-week-old female CD-1 mice administered 100 μg of indicated Fc-fusion protein prior to subcutaneous challenge with EEEV FL93–939. The scoring system is described in STAR Methods. (**D)** Survival of mice treated as in (**A**) following aerosol challenge with EEEV FL93–939. Data in **A-D** are pooled from two independent experiments with n = 10 mice per experimental group. (**A**) log-rank test with Bonferroni correction; (**D**) log-rank test: **p < 0.01, ****p < 0.0001.

## DISCUSSION

Although alphaviruses share similar virion structures, the glycoproteins vary substantially in sequence (∼45 to 90% identity for E1 and ∼40 to 80% identity for E2 across all serocomplexes),^53^ which likely contributes to their differential receptor usage. Previously characterized alphavirus receptors include MXRA8, which is used by several members of the Semliki Forest (SF) serocomplex (*e.g*., CHIKV, RRV, MAYV, and ONNV, but notably not by SFV),^37^ and LDLRAD3, which is used only by VEEV serocomplex members.^40^ Despite dissimilarity in protein topology, MXRA8 (Ig-like domains) and LDLRAD3 (LA domains) engage their respective viruses within an analogous cleft formed by neighboring E1/E2 glycoprotein heterodimers.^41–44^ Recently, VLDLR and the related ApoER2 molecule were reported to bind and facilitate infection of EEEV, SFV, and to a lesser extent, SINV.^45^ While SFV was unexpectedly shown to bind LA3 via the domain III tail of E1,^46^ the binding mode of EEEV and VLDLR remained unclear.

Our experiments show that EEEV uses multiple distinct sites on the E2 glycoprotein to mediate efficient VLDLR-dependent infection (**Fig S8**). Whereas 2 of the 3 identified sites in E2 are conserved in all deposited EEEV structural polyprotein sequences present in Genbank, the PE–6 strain and some other related strains (EEEV A61-1K and Madariaga virus BeAr348998) have a rare polymorphism (E206K) that enables LA domain binding to a third site on the E2-B domain. In addition, we show that the five VLDLR LA domains (LA1, LA2, LA3, LA5, and LA6) that encode a conserved Trp all can participate in the binding to EEEV. Through a series of cryo-EM reconstructions and functional studies with different domain-containing fragments of VLDLR, we observed promiscuity in LA domain interactions with each of the three EEEV binding sites: (a) EEEV engages both VLDLR LA1 and LA2, with LA1 bound within the cleft near the E1 fusion loop and LA2 positioned atop a neighboring site on the A domain of E2; (b) LA3, LA5, and LA6 all can bind to the E2 B domain site when the K206 residue is present in EEEV; (c) our cryo-reconstruction of LA(3–8) show that domain(s) other than LA1 can bind in the cleft; and (d) while we were unable to obtain direct structural evidence for a non-LA2 domain binding at the A domain site, infection or neutralization of FL93–939 (which lacks the B domain site) with an LA(1–6) featuring an inactivating LA2 W89 mutation strongly suggests that an additional LA domain (*e.g*., likely LA3 or LA6) can bind the E2-A domain site. Although we did not observe conformational changes upon LA domain binding at any of the sites in EEEV, the potential effects of the different VLDLR binding modes on downstream processes of viral entry warrant further study.

The PE–6 strain, which is a component of the trivalent VEEV/EEEV/WEEV VLP vaccine currently in advanced clinical trials,^54^ features E2-K206, which allows EEEV to engage LA domains on the side of the E2 B domain. In contrast, other commonly used experimental EEEV strains (*e.g*., FL93–939 and FL91–469) possess E2-E206 and do not use their E2 B domain to engage VLDLR. Thus, the third VLDLR binding site observed for EEEV PE–6 appears to be an exception to the norm. While the possibility of sampling bias in EEEV isolates remains, nearly all other 692 EEEV sequences we analyzed possess E2-E206. The E2-K206 residue might be a tissue culture adaptation, since the sequence of the PE–6 strain was obtained after indeterminant passage in eggs, mice, and guinea pigs.^55^ Notwithstanding this point, one other EEEV isolate (A61-1K) and one sequenced Madariaga (EEE serocomplex) isolate also harbor E2-K206, although their usage of VLDLR as a receptor and the contribution of the E2 B domain warrants further study. Regardless, our structural and functional data show how viruses can acquire multiple receptor-binding sites to co-opt repeat domains present in host receptors for high avidity interactions.

Whereas SFV binds the same VLDLR LA domains as EEEV (LA1, LA2, LA3, LA5, and/or LA6) using a single site on domain III of E1 (located around the 5- and 2-fold axes of symmetry), EEEV engages VLDLR LA(1–2) more like CHIKV/MXRA8 and VEEV/LDLRAD3, as it dominantly interacts with sites on E2 and the E1 fusion loop within the E1/E2 cleft. This differential engagement of LA domains through different envelope protein sites helps to explain how distantly related alphaviruses can bind the same host receptor. LDL-receptor family members also have been implicated in entry of unrelated viral families; in every case that has been structurally characterized, which includes vesicular stomatitis virus (VSV)/LDLR,^56^ human rhinovirus 2 (HRV2)/VLDLR,^57^ VEEV/LDLRAD3,^43,44^ and SFV/VLDLR^46^, the virus has used lysine or arginine residues to target conserved aspartate, glutamate, and tryptophan residues in the calcium-coordination site of the LA repeat, suggesting convergence of unrelated or distantly related viruses toward a shared receptor engagement strategy. However, in the case of LA domains engaged in the E1/E2 cleft (*e.g.*, LDLRAD3 LA1-VEEV and VLDLR LA1-EEEV), other surfaces of the LA domain appear to augment specificity of the interaction. Notably, SFV E2 lacks the lysine (E2-K156) critical for VLDLR LA1 binding by EEEV (**Fig S6**), and EEEV E1 reciprocally lacks the key lysine (E2-K345) used by SFV to target LA3 (**Fig S5**),^46^ potentially explaining their disparate modes of receptor engagement.

In contrast to the LDLRAD3-LA1/VEEV interaction, EEEV requires concurrent use of at least two LA domains of VLDLR for an avid interaction and efficient infection. While a higher affinity, single-domain receptor could still exist for EEEV, analogous to LDLRAD3-LA1/VEEV, the use of multiple LA domains could afford greater versatility in receptor binding. Rather than co-evolving with a single protein to achieve high-affinity binding mediated by a large binding interface, some alphavirus structural proteins may have evolved toward more flexible domain usage, engaging host receptors through a conserved structural element (*e.g.*, the calcium-coordination site of the LA domain) with only a small interface, but at multiple sites for high avidity binding. Targeting a minimal, conserved feature also may enable interactions with related receptors in the same host or across orthologs in different species. Indeed, human *LRP8* (ApoER2) and the *Aedes aegypti* and *Aedes albopictus* VLDLR orthologs can support infection of EEEV.^45^ However, structure-based sequence alignment of VLDLR orthologs from species that do (*Aedes* species and human) or do not (horse, avian, and *C. elegans*) support EEEV infection^45^ does not readily explain the differential receptor usage across evolution, as critical interacting residues within the calcium-coordination site generally are conserved across species (**Fig S9**). This suggests that other sequence or structural features must dictate LA domain geometry and receptor utilization. It is also possible that EEEV engages *Aedes* species VLDLR orthologs in a different manner or at sites not used for human VLDLR, further complicating sequence-based analyses.

### Limitations of the study

We acknowledge limitations of our study. (1) Due to biosafety considerations, many of the structural and functional experiments were performed with EEEV VLPs or attenuated chimeric viruses (*e.g*., SINV-EEEV) that produce virions displaying the EEEV structural proteins; (2) While we generated a VLDLR LA(1–2)-Fc decoy that protected against subcutaneous or aerosol challenge with highly pathogenic EEEV, we did not test its post-exposure activity; (3) Some of our cryo-EM reconstructions are of moderate resolution, limiting our ability to make claims of specific molecular interactions between EEEV and VLDLR; and (4) We did not test VLDLR LA domain decoy molecules against the passaged EEEV–PE6 strain *in vivo* to assess whether engagement of the E2 domain B binding site would confer greater protection.

In summary, the alphavirus-receptor complexes described here and elsewhere together demonstrate how modes of engagement between receptor and virus can defy expectations, as the notion of a sole “receptor-binding domain” may be oversimplified. Different homologs and/or orthologs of LA domains or other types of repeat domains (*e.g.,* Ig-like) could be engaged at spatially distinct sites or in differing orientations by the same or related viruses. A more complete understanding of virus-receptor biology and entry will require characterization of more receptors in relevant hosts to determine how viruses evolve to infect cells and adapt to new species.

## Acknowledgements

We thank Michael Rau, Brock Summers, Katherine Basore, and James Fitzpatrick at the Washington University Center for Cellular Imaging (WUCCI) for cryo-EM sample preparation and data acquisition. We thank Camille Carmona for technical assistance. We acknowledge K. Carlton and J. Mascola from the Vaccine Research Center of the National Institutes of Allergy and Infectious Diseases (NIH) for a gift of the EEEV PE–6 VLPs. The studies were supported by NIH grants R01 AI141436 and contract AI201800001 (to M.S.D.), U19 AI142790 (to M.S.D and D.H.F.), NIAID contract HHSN272201700060C (to D.H.F.), R01AI095436 (to W.B.K.), and DTRA MCDC2103-01 (to W.B.K. and M.S.D.)

## Author Contributions

L.J.A., S.R. and M.S.D. designed the study. L.J.A. performed cryo-EM reconstruction and model building and generated all structural representations. S.R. and L.J.A. generated Fc-fusion proteins. S.R. generated VLPs and performed all cell culture and mutagenesis experiments. L.J.A. performed affinity analyses. H.M. provided critical reagents and experimental support. T.G., D.S.R, and W.K. performed and supervised *in vivo* studies. D.H.F. supervised and analyzed the cryo-EM and binding studies. L.J.A., S.R. and M.S.D. wrote the initial draft with all other authors providing comments.

## SUPPLEMENTAL FIGURE LEGENDS

**Figure S1. Cryo-EM methodology. Related to Figure 1.** Cryo-EM data processing steps are shown for EEEV VLPs apo, in complex with full-length VLDLR, or in complex with VLDLR LA(1–2).

**Figure S2. Cryo-EM quality control. Related to Figure 1.** (**A-C**) Local resolution estimates (*left*) and gold-standard FSC plots (*right*) of EEEV asymmetric unit apo (**A**), in complex with full-length VLDLR (**B**), or in complex with VLDLR LA(1–2) (**C**). Resolutions were estimated in cryoSPARC using a 0.143 FSC cutoff. (**D**) Example model fits at EEEV-VLDLR interfaces, with experimental cryo-EM densities shown as a mesh.

**Figure S3. Defining VLDLR LA domains that support EEEV PE–6 binding and infection. Related to Figure 2.** (**A**) Phylogenetic tree generated using the sequences of the structural genes E1 and E2 of the indicated alphaviruses. Colored lines indicate known receptor usage by the corresponding virus. (**B**) Flow cytometry plots of K562 cells stained with the indicated mAbs (*left*) and representative flow plot of GFP expression in K562 cells following SINV-EEEV-GFP expression (*right*). (**C**) Flow cytometry histograms showing expression of the indicated Flag-tagged VLDLR constructs in K562 cells. (**D**) Schematic of SINV chimeric reporter viruses in which the structural genes of SINV have been replaced with those of the indicated alphaviruses in addition to GFP. (**E**) Infection of K562 cells expressing the indicated constructs by SINV-VEEV-GFP (*left*) and SINV-EEEV-GFP (*right*) as assessed by GFP expressing using flow cytometry. Data are pooled from two to four independent experiments. (**F**) Flow cytometry histograms showing expression of the indicated Flag-tagged truncated VLDLR constructs in K562 cells. (**G**) Alignment of the 8 LA domains of VLDLR and LDLRAD3 LA1. Filled or open arrowheads respectively indicate residues that coordinate calcium by side chain or main chain carbonyl. (**H**) Flow cytometry plots of the indicated single LA domain transduced K562 cells showing expression of the Flag tag in control K562 and K562-VLDLR cells. Flow cytometry histograms in **B, C, F, and H** are representative of two independent experiments. Data in **E** are pooled from two independent experiments.

**Figure S4. Characterization of LA domain binding of VLDLR by EEEV PE–6. Related to Figure 2.** (**A**) Diagram of BLI experiments. Fc-fusion proteins are captured with anti-human Fc biosensors followed by incubation with VLPs (left schematic, “receptor immobilized”). VLPs are captured by anti-mouse mAbs (EEEV-3) followed by incubation with Fc-fusion protects (right schematic, “receptor in solution”). (**B**) BLI of Fc-fusion proteins following incubation with VLPs. Representative sensor traces are shown after dipping into wells containing 20 μg/mL of EEEV (left panel) or VEEV VLP (right panel). Data are representative of two independent experiments. (**C, E, F, G**) Flow cytometry histograms showing expression of the indicated Flag-tagged chimeric VLDLR constructs in K562 cells. (**D**) Quantification of GFP^+^ cells (%) 24 h after SINV-VEEV-GFP TrD infection of K562 cells expressing indicated VLDLR(WA) constructs from (**C**). (**H)** Quantification of GFP^+^ cells (%) 24 h after SINV-SFV4-GFP infection of K562 cells expressing WT VLDLR or LA(2–3)ΔLBD. Data are pooled from 4 independent experiments. (**I)** Flag expression of indicated VLA(1–2)ΔLBD mutants as assessed by flow cytometry. Statistical analysis (**D, H**): n.s, not significant, **** p < 0.0001; student’s t-test.

**Figure S5. Characterization of EEEV FL93–939 binding and infection by VLDLR. Related to Figure 5.** (**A**) Dose-response curves of neutralization by the indicated Fc-fusion proteins against SINV-EEEV FL93–939-GFP. (**B)** Steady-state BLI curves of monovalent LA(1–6) bound to indicated EEEV FL93–939 VLP coated biosensors. Data are pooled from three independent experiments. (**C**) BLI of Fc-fusions following incubation with VLPs. Representative sensor traces are shown after dipping into wells containing 20 μg/mL of EEEV FL93–939 VLPs. (**D-E**) SINV-EEEV-GFP FL93–939 infection of K562 cells transduced with variants of VLDLR (WA) in which the single (**D**) or two (**E**) LA domains have been reverted to Trp. (**F-H**) Scheme of LA(1–6)-Fc fusion protein and relevant Trp residues annotated (**F**). Neutralization by the indicated Trp variants of LA(1–6)-Fc proteins against SINV-EEEV-GFP PE–6 (**G**) or SINV-EEEV-GFP FL93–939 (**H**) in 293T cells. (**I)** Binding of LA(1–6mut2)-Fc to captured wild-type (WT) and mutant EEEV FL93–939 VLPs. Biosensors were coated with WT or mutant VLPs followed by incubation with 25 nM of LA(1–6mut2)-Fc for 300 sec. Binding was calculated as percent signal (R_max_) relative to WT VLPs. Data in (**A, G, H**) are pooled from two to three independent experiments. Data in (**E, F, I**) are pooled from three independent experiments: *p < 0.05, ****p < 0.0001, n.s. not significant; one-way ANOVA with Dunnett’s post-test).

**Figure S6. Alphavirus E1 multiple sequence alignment with receptor contacts. Related to Figure 6.** Structural alignment of E1 proteins from EEEV PE–6 (GenBank L37662.1), EEEV FL93–939 (EF151502.1), VEEV (strain TC-83, AAB02517.1), SFV (strain 4, AKC01668.1), and CHIKV (strain 37997, AAU43881.1) generated with PROMALS3D^58^ and visualized with ESPript 3.^59^ Receptor contacts (determined by PISA) are shown below the alignment in orange (EEEV/VLDLR; transparent orange for PE–6-specific contacts), yellow (VEEV/LDLRAD3), purple (CHIKV/MXRA8), or green (SFV/VLDLR), delineated as wrapped, intraspike, interspike (CHIKV/MXRA8 only), or vertex (SFV/VLDLR only). Blue boxes highlight electropositive residues on a given virus known to form salt bridges with the calcium-coordination site of an LA domain receptor.

**Figure S7. Alphavirus E2 multiple sequence alignment with receptor contacts. Related to Figure 6.** Structural alignment of E2 proteins from EEEV PE–6 (Genbank L37662.1), EEEV FL93–939 (EF151502.1) VEEV (strain TC–83, AAB02517.1), SFV (strain 4, AKC01668.1), and CHIKV (strain 37997, AAU43881.1) generated with PROMALS3D^58^ and visualized with ESPript 3.^59^ Receptor contacts (determined by PISA) are shown below the alignment in orange (EEEV/VLDLR; transparent orange for PE–6-specific contacts), yellow (VEEV/LDLRAD3), or purple (CHIKV/MXRA8), delineated as wrapped, intraspike, or interspike (CHIKV/MXRA8 only). Blue boxes highlight electropositive residues on a given virus known to form salt bridges with the calcium-coordination site of an LA domain receptor.

**Figure S8. Differential VLDLR LA domain usage by distinct binding sites on EEEV**. **Related to Figures 2-5. A**. Schematic representation of VLDLR LA domain usage at the different receptor-binding sites on EEEV (E1/E2 cleft, E2 A domain, and E2 B domain). The cleft and E2 A sites are conserved in all EEEV strains whereas the E2 B domain binding site is present in the few strains featuring residue E2-206K (e.g., EEEV PE–6). Arrows from LA domains indicate which sites on EEEV are bound, respectively, with solid lines indicating interactions observed structurally, and thicker lines indicating interactions that are of higher affinity. **B**. Predicted EEEV-VLDLR binding modes are depicted on the virion surface.

**Figure S9. Sequence alignment of VLDLR orthologs, Related to Figures 5-6.** Multiple sequence alignment of *Homo sapiens* (human, GenBank NP_003374.3), *Mus musculus* (mouse, NP_038731.2), *Equus caballus* (horse, XP_023483037.1), *Sturnus vulgaris* (avian, XP_014736085.1), *Aedes aegypti* (mosquito, AEY84776.1), *Aedes albopictus* (mosquito, JAC13440.1), and *Caenorhabditis elegans* (nematode, NP_872023.2) VLDLR orthologs. Structural homology guided alignment was performed via cysteine barcoding^60^ followed by alignment via PROMALS3D,^58^ visualized with ESPript 3.^59^ LA domains are annotated below the alignment. Predicted EEEV contacts are designated by large (close contacts) or small (other contacts) dots, with strain-specific contacts (in LA6) colored lightly.

## SUPPLEMENTAL TABLE TITLES

**Supplemental Table 1. Cryo-EM data collection, processing, and model refinement statistics. Related to Figures 4 and 5.**

**Supplemental Table 2. Residues comprising the EEEV/VLDLR interface. Related to Figures 4 and 5.** Contact residues were determined by PISA. Close contacts (≤ 3.9 Å) conserved at all binding sites in the asymmetric unit are underlined. Strain (PE–6)-specific contacts are italicized.

**Supplemental Table 3. Sequence alignment of EEEV sequences in Genbank, Related to Figure 5**

## STAR METHODS

### RESOURCE AVAILABILITY

#### Lead contact

Further information and requests for resources and reagents should be directed to the Lead Contact, Michael S. Diamond.

#### Materials availability

All requests for resources and reagents should be directed to the Lead Contact author. This includes viruses, proteins, and cells. All reagents will be made available on request after completion of a Materials Transfer Agreement (MTA).

#### Data and code availability

All data supporting the findings of this study are available within the paper and are available from the corresponding author upon request. This paper does not include original code. All structures have been deposited in the PDB and EMDB databases (PDB: 8SNT, 8SNU; EMDB: 17237, 40634, 40635, 40636).

### EXPERIMENTAL MODEL AND SUBJECT DETAILS

#### Cells

293T (ATCC, CRL-3216). Vero (ATCC, CCL-81), and BHK21 (ATCC, CCL-10) cells were maintained in high-glucose DMEM supplemented with 10% FBS, Gluta-MAX, 10 mM HEPES, non-essential amino acids, and penicillin-streptomycin. K562 (ATCC, CCL-243 cells were maintained in RPMI-1640 (Thermo Fisher) supplemented with 10% FBS, Gluta-MAX and 10mM HEPES.

#### Viruses

SINV-EEEV-EGFP PE–6 (and mutants) and SINV-SFV4-EGFP were generated by replacement of structural genes of SINV-WEEV-EGFP CBA87 with EEEV PE–6 and SFV4 structural genes, respectively, by PCR and Gibson assembly with EGFP expressed a structural protein fusion as described previously.^61^ The infectious cDNA clones were digested with XhoI, and the linearized vector purified with Monarch PCR & DNA Clean up Kit (New England BioLabs). One μg of linearized vector was used to generate RNA with a HiScribe SP6 RNA synthesis kit (New England BioLabs) followed by purification with Monarch RNA clean up kit. Four μgs of RNA were transfected into BHK21 cells with a GenePulser Xcell electroporator (Bio-Rad). The supernatant was harvested as the P0 stock 48 h later. The virus was passaged one additional time on BHK21 cells and titered on Vero cells. SINV-VEEV-EGFP TrD and SINV- EEEV-GFP FL93–939 has been described previously. EEEV FL93–939 was produced from a cDNA clone (a gift from S. Weaver, UTMB Galveston) as described previously.^32^

#### Mouse studies

All animal procedures performed at the University of Pittsburgh were carried out under approval of the Institutional Animal Care and Use Committee of the University of Pittsburgh in protocols 15066059 and 18073259. Animal care and use were performed in accordance with the recommendations in the Guide for the Care and Use of Laboratory Animals of the National Research Council. Approved euthanasia criteria were based on weight loss and morbidity. Virus inoculations were performed under anesthesia that was induced and maintained either with inhaled isoflurane or ketamine hydrochloride and xylazine, and all efforts were made to minimize animal suffering. CD-1 mice were purchased commercially (Jackson Laboratories). Subcutaneous infections with cDNA clone-derived EEEV FL93–939 were in the left rear footpad and aerosol infections were performed as previously described.^62^

### METHOD DETAILS

#### Ectopic expression of VLDLR constructs

cDNA encoding VLDLR (GenBank NP_003374.3) residues 28-873 was subcloned into a pLV-IRES-puromycin lentiviral vector with an N-terminal Flag tag, downstream of a human β2m signal sequence. VLDLR chimeras and mutants were cloned into the pLV-VLDLR-IRES-puro or by Gibson assembly with Geneblocks (IDT) and verified by Sanger sequencing or were designed and synthesized by Twist Biosciences (San Francisco, CA), into the pSFFV-IRES-puromycin backbone. Lentivirus was harvested 48 h post-transfection of HEK293T cells with donor vector, psPAX2 (Addgene #12260), and pMD2.G (Addgene#12259) in a 2:2:1 ratio using Mirus LT-1 reagent according to the manufacturer’s recommended instructions. We ‘spinoculated’ (800 x *g* for 25 min) K562 cells with the lentiviral supernatants and exchanged cells into fresh media after 24 hours. Transduced cells were selected after 48 h of incubation with puromycin (2 μg/mL, Invivogen) and 7 days of culture before use in experiments. To verify Flag-tag expression after selection, cells were incubated with anti-Flag-Alexa Flour 647 (15009S, Cell Signaling) at a 1:200 dilution in FACS buffer (1x PBS + 0.1% BSA + 2 mM EDTA + 0.05% NaN_3-_) for 30 min at 4°C. Following washing steps, cells were analyzed on an iQue3 flow cytometer. For some experiments, cells stained with anti-Flag-A647 were sorted for either low or high expression on a MACSQuant Tyto instrument (Miltenyi Biotec).

#### Cell infection experiments

Transduced K562 cells were inoculated with SINV–EGFP chimeric viruses at a multiplicity of infection (MOI) of 2 for 24 h. To assess the neutralization capacity of Fc-fusion proteins, virus was incubated with the VLDLR Fc-fusion proteins for 1 h at 37°C before inoculation of 293T cells. Cells were harvested 16 h post-infection for flow cytometric analysis. Cells were subjected to flow cytometry in the presence of 4’,6-diamidino-2-phenylindole (DAPI, 1μg/mL) using an iQue3 (Sartorius) and analyzed with Forecyte (Sartorius) software.

#### Fc-fusion proteins

The ligand binding domain (LA1-8) of human VLDLR (residues 28– 355, GenBank NP_003374.3) was subcloned into a pTwist CMV β-globin expression vector downstream of a mouse IgH signal sequence encoding human IgG1 Fc separated by a GGGSGGS linker with or without an HRV 3C protease cleavage site. The individual and truncated Fc-fusion constructs were generated in a similar fashion: LA1 (31-69), LA2 (70-110), LA3 (111-151), LA4(152-190), LA5(191-231), LA6(237-275), LA7(276-314), LA(1–2) (31-110), LA(2–3) (70-151), LA(3–4) (111-190), LA(4–5) (152-231), LA(5–6) (191-275), LA(1–3) (31-151), LA(1–4) (1-190), LA(1–5) (1-231), LA(1–6) (1-275). To express proteins, constructs were co-transfected with human LRPAP (RAP) chaperone protein (NM_002337.4, residues 1– 353) at a (4:1 ratio) with Expifectamine 293 reagent into Expi293 cells (Thermo Fisher) according to the manufacturer’s instructions. Supernatants were harvested four days post-transfection, centrifuged, filtered, and Fc-fusion proteins were bound to Protein A Sepharose (Thermo) in a gravity flow column. The column was washed with 25 column volumes of 1x TBS (20 mM Tris pH 8.0, 150 mM NaCl), 50 column volumes of high-salt buffer (20 mM Tris pH 8.0, 500 mM NaCl, 10 mM EDTA) to strip LRPAP, followed by 25 column volumes of 1x TBS + 10 mM CaCl_2_. Proteins were eluted with Pierce™ Gentle Ag/Ab Elution Buffer, pH 6.6 (Thermo Fisher) and desalted into 1x TBS (20 mM Tris pH 8.0, 150 mM NaCl) with 2 mM CaCl_2_ using a PD-10 column (Cytiva). Depending on the yield, protein was concentrated with an Amicon 10 kDa centrifugal filter (Millipore). Protein purity was assessed by non-reducing SDS-PAGE followed by staining with SimpliSafe Coomasie reagent (Thermo). Gels were imaged with an iBright 1500 instrument (Thermo Fisher).

#### VLPs

EEEV PE–6 and VEEV TC-83 VLPs were a gift from K. Carlton and J. Mascola, Vaccine Research Center of the National Institutes of Allergy and Infectious Diseases).^63^

#### Domain mapping and quantitative BLI

All domain mapping and quantitative affinity experiments were performed on a GatorPlus BLI and analyzed using on-board software (GatorBio) or BiaEvaluation Software (Biacore). Unless otherwise noted, all experiments were performed with 1x PBS supplemented with 1% BSA and 2 mM CaCl_2_ (Running buffer). To test the avidity of receptor constructs clustered in the solid phase, VLDLR-Fc fusion constructs were immobilized on anti-human IgG Fc biosensors (GatorBio #160003) for 120 sec, washed in Running buffer for 30 sec, then submerged in EEEV or SFV VLPs at a nominal concentration of ∼20 μg/mL for 360 sec to assess binding. To evaluate binding of VLDLR constructs in solution, anti-mouse IgG Fc biosensors (GatorBio #160004) were incubated ∼20 μg/mL of mEEEV-3^50^ for 120 sec then, after washing in Running buffer for 30 sec, EEEV VLPs were captured at a nominal concentration of ∼20 μg/mL for 240 sec. For qualitative domain mapping, VLP-coated biosensors then were dipped into wells containing 1 μM of each VLDLR Fc-fusion construct; for quantitative kinetic experiments, VLP-coated biosensors were submerged into in the indicated concentrations of monovalent VLDLR fragmentscleaved from the Fc using GlySERIAS (Genovis #A0-GS6) or Pierce™ HRV 3C protease (Thermo Fisher). Steady state (equilibrium) affinity was determined via on-board on-board GatorOne Software (v2.7, GatorBio). Experiments were performed in replicate (qualitative) or triplicate (quantitative).

#### Mutant VLP generation and assessment of VLDLR-Fc binding

To produce mutant EEEV VLPs, we utilized a previously generated EEEV E2 alanine-scanned library or E1 mutants of a pCAGGS vector encoding the structural proteins of EEEV (KR780-2). Select E2 mutants were transfected into Expi293 cells and supernatants were harvested after 2 days and centrifuged to remove cells and debris. VLPs were captured from the crude supernatant with mEEEV-3 coated anti-mouse IgG biosensors. We then saturated non-specific binding sites on the anti-mouse IgG pins by incubating with 20 μg/mL of MAY-117 (an isotype control mAb). To assess and quantify binding to LA(1–2)-Fc, the EEEV VLP coated sensor tips were dipped into 1000 nM of LA(1–2)-Fc or 10 μg/mL of EEEV-3. The percent binding was quantified as the BLI signal of LA(1–2)-Fc relative to EEEV-3, with mutant VLPs normalized to that of WT VLPs.

#### Mouse infections

Five-week old CD-1 female mice were injected intraperitoneally with a single 100 mg (∼5 mg/kg) dose of VLDLR LA(1–2)-Fc or LDLRAD3 LA1-Fc or PBS (in 200 ml PBS volume), followed 6 h later by subcutaneous inoculation with 10^3^ plaque-forming units (PFU) of EEEV FL93–939 in the left rear footpad. Aerosol exposures were performed as previously described^62^ using the AeroMP exposure system (Biaera Technologies) inside a biological safety cabinet class III with target dose of 5 x 10^2^ PFU. Mice were monitored once or twice daily for weight loss, morbidity, and mortality through 14 days post infection. Clinical signs were assigned by the following criteria: 0 - healthy; 1 - ruffled fur, mild behavioral changes; 2 - hunched posture, significant behavioral changes; 3 - seizures, ataxia, catatonia; 4 - recumbent moribundity; 5 - death. Mice scoring 3 or higher were immediately euthanized.

#### Cryo-EM sample preparation

EEEV VLPs were prepared at a nominal concentration of ∼ 0.7 mg/mL in PBS (pH 7.4), then incubated for 1 h on ice with 2-to-10-fold molar excess of full-length VLDLR (ACROBiosystems # VLR-H5227) or VLDLR fragments. Solutions of VLPs alone or with VLDLR were applied to glow-discharged lacey carbon grids (Ted Pella #01895-F) then flash-frozen in liquid ethane using a Vitrobot Mark IV (ThermoFisher Scientific).

#### Cryo-EM data collection

Grids were loaded into a Cs-corrected FEI Titan Krios 300kV microscope or an FEI Glacios 200kV scope, each equipped with a Falcon 4 direct electron detector, and then imaged at a nominal magnification of 59k× (Krios) or 120k× (Glacios), resulting in a calibrated pixel size of 1.081 Å (Krios) or 1.184 Å (Glacios). Movies were collected in EER format with a total dose of ∼36-40 e^-^/Å^2^/movie (∼4.5 e^-^/Å^2^/s over 8-9 sec acquisition).

#### Cryo-EM data processing

EER movies were binned into 36-40 fractions (∼1.0 e-/Å^2^/f) and then pre-processed via patch motion and patch CTF correction in cryoSPARC v3.1.0.^64^ Particles were selected using a template picker then cleaned via two- and three-dimensional classification. Whole VLPs with or without VLDLR were reconstructed via homogeneous non-uniform refinement with I1 symmetry imposed. We then performed symmetry expansion and extracted individual asymmetric units for focused three-dimensional classification without orientational sampling in Relion 3.1,^65^ and the class of highest resolution for each sample was subjected to local non-uniform refinement in cryoSPARC.

#### Model building

Starting models for EEEV structural proteins were adapted from a previous EEEV VLP cryo-EM structure (PDB: 6XO4),^47^ and VLDLR LA domains were modeled using AlphaFold2 implemented in ColabFold.^66,67^ These starting components were docked into the electron density maps and then refined iteratively using Coot v0.9.5,^68^ Isolde v1.1.0,^69^ Phenix v1.19,^70^ and Rosetta scripts.^71–73^ Proteins, Interfaces, Structures, and Assemblies (PISA)^51^ solvent exclusion analysis was used to identify contact residues and calculate buried surface area. Structures were visualized using UCSF ChimeraX.^74^

### QUANTIFICATION AND STATISTICAL ANALYSES

Statistical significance was assigned using Prism Version 8.0 (GraphPad) when *P* < 0.05. Statistical analysis of viral infection levels was determined by one-way ANOVA with Dunnett’s post-test or student’s t test. Statistical analysis of *in vivo* experiments was determined by Kaplan-Meier survival curve analysis. The statistical tests, number of independent experiments, and number of experimental replicates are indicated in the Figure legends.

