## Supplementary Table S1 for "Structural and functional basis of VLDLR receptor usage by Eastern equine encephalitis virus"

### Table S1. Cryo-EM data collection, processing, and model refinement statistics, Related to Figures 4 and 5

| **Sample** | EEEV<br>VLP | EEEV<br>VLP<br>PE–6<br>+<br>full-<br>length<br>VLDLR | EEEV<br>VLP<br>PE–6<br>+<br>VLDLR<br>LA(1–2) | EEEV<br>VLP<br>PE–6<br>+<br>VLDLR<br>LA(1–3) | EEEV<br>VLP<br>PE–6<br>+<br>VLDLR<br>LA(1–5)<br>W132A | EEEV<br>VLP<br>PE–6<br>+<br>VLDLR<br>LA(1–6)<br>W132A<br>W210A | EEEV<br>VLP<br>PE–6<br>+<br>VLDLR<br>LBD<br>W132A<br>W210A<br>W256A | EEEV<br>VLP<br>PE–6<br>+<br>VLDLR<br>LA(3–8) | EEEV<br>VLP<br>PE–6<br>+<br>VLDLR<br>LA(1–6)<br>W89A | EEEV<br>VLP<br>FL93–939<br>+<br>VLDLR<br>LA(1–6) |
|---|---|---|---|---|---|---|---|---|---|---|
| EMDB | **VLP:**<br>42187<br>**ASU:**<br>42188 | **VLP:**<br>42189<br>**ASU:**<br>42190 | **VLP:**<br>42191<br>**ASU:**<br>42192 | **VLP:**<br>42193<br>**ASU:**<br>42194 | **VLP:**<br>42195<br>**ASU:**<br>42196 | **VLP:**<br>42197<br>**ASU:**<br>42198 | **VLP:**<br>42199<br>**ASU:**<br>42200 | **VLP:**<br>42201 | **VLP:**<br>42202<br>**ASU:**<br>42203 | **VLP:**<br>42212 |
| PDB | **ASU:**<br>8UFA | **ASU:**<br>8UFB | **ASU:**<br>8UFC | | | | | | | |
| **Data collection** | | | | | | | | | | |
| Voltage (kV) | 300 | 300 | 300 | 300 | 300 | 300 | 200 | 300 | 300 | 300 |
| Magnification | 59,000× | 59,000× | 59,000× | 59,000× | 59,000× | 59,000× | 120,000× | 59,000× | 59,000× | 59,000× |
| Exposure (e-/Å2) | 37.93 | 37.90 | 37.90 | 37.90 | 44.40 | 44.40 | 39.90 | 44.57 | 44.40 | 44.13 |
| Defocus range (μm) | 0.7-2.2 | 0.7-2.2 | 0.7-2.2 | 0.7-2.2 | 0.7-2.2 | 0.7-2.2 | 0.5-1.9 | 0.7-2.2 | 0.7-2.2 | 0.7-2.2 |
| Pixel size (Å/pixel) | 1.081 | 1.081 | 1.081 | 1.081 | 1.081 | 1.081 | 1.184 | 1.081 | 1.081 | 1.081 |
| **Data processing** | | | | | | | | | | |
| Final particles (no.) | **VLP:**<br>41,227<br>**ASU:**<br>1,038,056 | **VLP:**<br>13,998<br>**ASU:**<br>391,827 | **VLP:**<br>19,222<br>**ASU:**<br>459,518 | **VLP:**<br>19,321<br>**ASU:**<br>491,574 | **VLP:**<br>1,465<br>**ASU:**<br>55,592 | **VLP:**<br>4,724<br>**ASU:**<br>144,008 | **VLP:**<br>2,813<br>**ASU:**<br>84,264 | **VLP:**<br>13,331 | **VLP:**<br>1,057<br>**ASU:**<br>41,163 | **VLP:**<br>9,669 |
| Resolution (Å) | **VLP:**<br>3.78<br>**ASU:**<br>2.86 | **VLP:**<br>4.75<br>**ASU:**<br>3.89 | **VLP:**<br>4.09<br>**ASU:**<br>3.09 | **VLP:**<br>4.21<br>**ASU:**<br>3.24 | **VLP:**<br>4.94<br>**ASU:**<br>3.71 | **VLP:**<br>4.05<br>**ASU:**<br>2.98 | **VLP:**<br>6.03<br>**ASU:**<br>4.75 | **VLP:**<br>3.66 | **VLP:**<br>5.72<br>**ASU:**<br>4.26 | **VLP:**<br>4.72 |
| FSC threshold | 0.143 | 0.143 | 0.143 | 0.143 | 0.143 | 0.143 | 0.143 | 0.143 | 0.143 | 0.143 |
| **Model refinement** | | | | | | | | | | |
| Initial models | 6XO4 | 6XO4 | 6XO4 | | | | | | | |
| Model resolution (Å) | 2.8/3.1 | 3.8/4.3 | 3.0/3.4 | | | | | | | |
| FSC threshold | 0.143/0.5 | 0.134/0.5 | 0.143/0.5 | | | | | | | |
| Model composition | | | | | | | | | | |
| Non-hydrogen atoms | 31,260 | 34,660 | 33,624 | | | | | | | |
| Residues | 4,024 | 4,476 | 4,336 | | | | | | | |
| Ligands (glycans) | 8 | 8 | 8 | | | | | | | |
| Bonds (RMSD) | | | | | | | | | | |
| length (Å) | 0.003 | 0.003 | 0.002 | | | | | | | |
| Angles (°) | 0.679 | 0.566 | 0.648 | | | | | | | |
| Validation | | | | | | | | | | |
| Molprobity score | 0.83 | 1.04 | 1.00 | | | | | | | |
| Clash score | 1.16 | 1.76 | 1.78 | | | | | | | |
| Rotamer outliers (%) | 0.00 | 0.00 | 0.00 | | | | | | | |
| Ramachandran | | | | | | | | | | |
| Favored (%) | 98.05 | 97.45 | 97.70 | | | | | | | |
| Allowed (%) | 1.95 | 2.55 | 2.30 | | | | | | | |
| Outliers (%) | 0.00 | 0.00 | 0.00 | | | | | | | |
