## Supplementary Table S2 for "Structural and functional basis of VLDLR receptor usage by Eastern equine encephalitis virus"

| **Table S2. List of residues comprising EEEV/VLDLR interface. Related to Figures 4 and 5.** | | | |
|---|---|---|---|
| **E1/E2 heterodimer** | **E1/E2 domain** | **E1/E2 residues** | **VLDLR residues** |
| wrapped | E2 A domain | R26, C27, D28 | **LA1:** R44 |
| | E2 B domain | H175, V220, K221, R224 | **LA1:** A31, K32, Q39, C40, T41, N42, G43, R44, D60 |
| | E1 fusion loop | Y85, F87, M88, Y89, G90, G91, A92, F95, D97, T98 | **LA1:** A31, K32, Q39, N42, G43, R44, C45, V59, D60 |
| | E1 domain II | G228, I229 | **LA1:** R44 |
| intraspike | E2 A domain | D1, L2, D3, T4, H5 K10, L11, K56, T57, D58, G59, V60, L62 | **LA1:** C52, D53, G54, D55, E64, C67<br>**Linker:** V68, K69, K70<br>**LA2:** C72, Q83, C84, V85, P86, R88, W89, D94, P95, D96, C97, E98 |
| | E2 B domain | S191, G192, N230 | **LA2:** R88, D92, G93, D94 |
| | E2 β-linker | T154, H155, K156, R157, A158, K231, K232, W233 | **LA1:** L49, W50, D53, D55, D57<br>**LA2:** R88, W89, D92, D94, P95, D96 |
| *PE-6 contacts* | *E2 B domain* | *Y197, Y198, K200, P202, D203, V204, R205, K206, G207, I208* | ***LA6:*** *E243, G249, E250, C251, H253, K255, W256, D259, D261, P262, D263, C264, K265* |
| | *E1 domain II* | *K61, C62, C63, G64* | ***LA6:*** *D259, G260, D261, P262* |
| | *E1 fusion loop* | *E99* | ***LA6:*** *P262* |
